# Investigating the Rescue Potential of Fecal Microbiota Transplants in Anorexia Nervosa using Antibiotic-treated Mice in a Split Group Cross-Over Study

**DOI:** 10.1101/2023.10.05.561007

**Authors:** Sina Maschek, Theresa Helena Østergaard, Lukasz Krych, Line Fisker Zachariassen, Dorte Bratbo Sørensen, Caroline M. Junker Mentzel, Axel Kornerup Hansen, Jan Magnus Sjögren, Kenneth Klingenberg Barfod

## Abstract

Anorexia nervosa (AN) is a complex and serious mental disorder, which may affect individuals of all ages and sex, but primarily affecting young women. The disease is characterized by a disturbed body image, restrictive eating behavior, and a lack of acknowledgment of low body weight. The underlying causes of AN remain largely unknown, and current treatment options are limited to psychotherapy and nutritional support. This paper investigates the impact of Fecal Microbiota Transplants (FMT) from AN patients on food intake, body weight, behavior, and gut microbiota in antibiotic-treated mice. Two rounds of FMT were performed using AN and control (CO) donors. During the second round of FMT, a subset of mice received FMT from a different donor type. This split-group cross-over design was chosen to demonstrate any recovery effect of FMT from a healthy donor. The first FMT, from AN donors, resulted in lower food intake in mice without affecting body weight. After FMT2, serum analysis revealed higher levels of appetite-influencing hormones (PYY and leptin) in mice receiving AN-GM. Gut microbiota analysis showed significant differences between AN and CO mice after FMT1, before cross-over. Specific bacterial genera and families *Ruminococcaceae, Lachnospiraceae,* and *Faecalibacterium* showed different abundances in AN and CO receiving mice. Behavioral tests showed decreased locomotor activity in AN mice after FMT1. Overall, the results suggest that AN-GM may contribute to altered food intake and appetite regulation, which can be ameliorated with FMT from a healthy donor potentially offering FMT as a supportive treatment for AN.

## Introduction

Anorexia nervosa (AN) is a complex and serious mental disorder, which may affect individuals of all ages and sex with the highest prevalence in young women (1, 2). Restrictive AN is characterized by a disturbed body image or lack of acknowledgement of low body weight and behavior that interferes with weight gain (3). AN is in some cases preceded by depression, trauma, gastrointestinal symptoms, an infection, or a combination hereof, however, in most cases, the etiology is largely unknown (4). Treatment options for AN are still limited to mainly psychotherapy and nutritional support and new avenues of approach to therapy are warranted.

There is growing evidence which supports the role of human gut microbiome (GM) in health, weight regulation and metabolic function, and, therefore, it may be hypothesized that GM plays a role in the etiology of AN. In this emerging field, several studies have now shown significant GM differences in patients with AN compared to healthy subjects, termed a dysbiosis (5–11). A microbiome dysbiosis is characterized by either loss of commensals, expansion of pathobionts, loss of microbial diversity, or combinations hereof (12). It has also been demonstrated that weight restoration and introduction to regular diets and therapy of AN patients does not rectify the dysbiosis of the AN-GM (13–15). Currently, two single case studies, and one protocol, have been published exploring the use of fecal matter transplantation (FMT) as a possible therapeutic approach to rectifying dysbiosis by transferring GM from healthy donors to AN patients (16–18).

In mice there are only a few studies investigating the potential transfer of AN phenotypical and behavioral traits by FMT from AN patients to germ-free (GF) mice in order to show causality. Fan et al. (2023) showed a larger initial decrease in body weight and a slower weight gain over time in mice receiving AN microbiota compared with mice that received healthy-control FMT. However, the study does not take food intake into consideration. Hata et al. (2019) reported reduced body weights and behavioral changes in offspring from GF mice who received fecal microbiota from AN patients (19). The AN-receiving mice showed anxiety-like and compulsive behaviors when undergoing open-field and marble-burying tests. In another study, Glenny et al. (2021) reported no relationship between AN-associated GM and changes in body weight gain in adult recipient GF female and male mice (20). All three studies make use of GF mice but differ with respect to inoculums, mouse strain and other methodologies.

Humanizing mouse models with human-to-mice FMT is an emerging concept with pros and cons that provides an ecologically relevant context to study the role of GM in health and disease. However, the major obstacles are low colonization rates from donor GM and immunological abnormalities observed in these mice (21). GF mice are usually the first choice for FMT studies, due to their immunological naivety towards living microbes and the absence of competition between the transplanted and the residual microbes present in non-GF models. However, the use of antibiotics (AB) to deplete native GM from mice should be considered when studying phenotype transfers already known to be associated with alterations in GM. AB depletion of GM allows for investigations of GM changes happening at certain life stages and in immune-competent mice (22).

In this study, our approach is to use human-to-mouse FMT to transfer AN phenotypic traits to adolescent AB treated immunocompetent female mice. We observe food intake per cage, body weight gain and behavioral characteristics over time. We apply a split-group cross-over design, as we hypothesize that FMT from a healthy control donor might partially rescue an AN-GM induced phenotype and vice versa. We correlate this to appetite related biomarkers in mouse serum at the point termination. We hypothesize that the AN phenotype induced by FMT from a patient into AB treated mice will be rescued by FMT from a healthy control donor. Additionally, we conduct behavioral open field tests, hypothesizing that mice with the AN phenotype would show less activity compared to CO mice.

## Results

### Fecal Microbiota Transplants from Anorexia Nervosa Patients lowers Food Intake without impacting Body Weights

*Body Weights:* No body weight differences were observed in mice receiving fecal microbiota transplants (FMT) from anorexic donors (AN mice) compared to mice receiving FMT from control donors (CO mice) following the first five weeks of fecal microbiota transplants (FMT1) (Figure 1 left). Likewise, there were no differences in body weights following the split cross-over and four weeks of the second FMT (FMT2) analyzed according to either the last received donor type (AN or CO) (data not shown) or the combined donor from both FMTs (ANAN, COAN, ANCO, COCO) (Figure 1 right).

**Figure 1.**
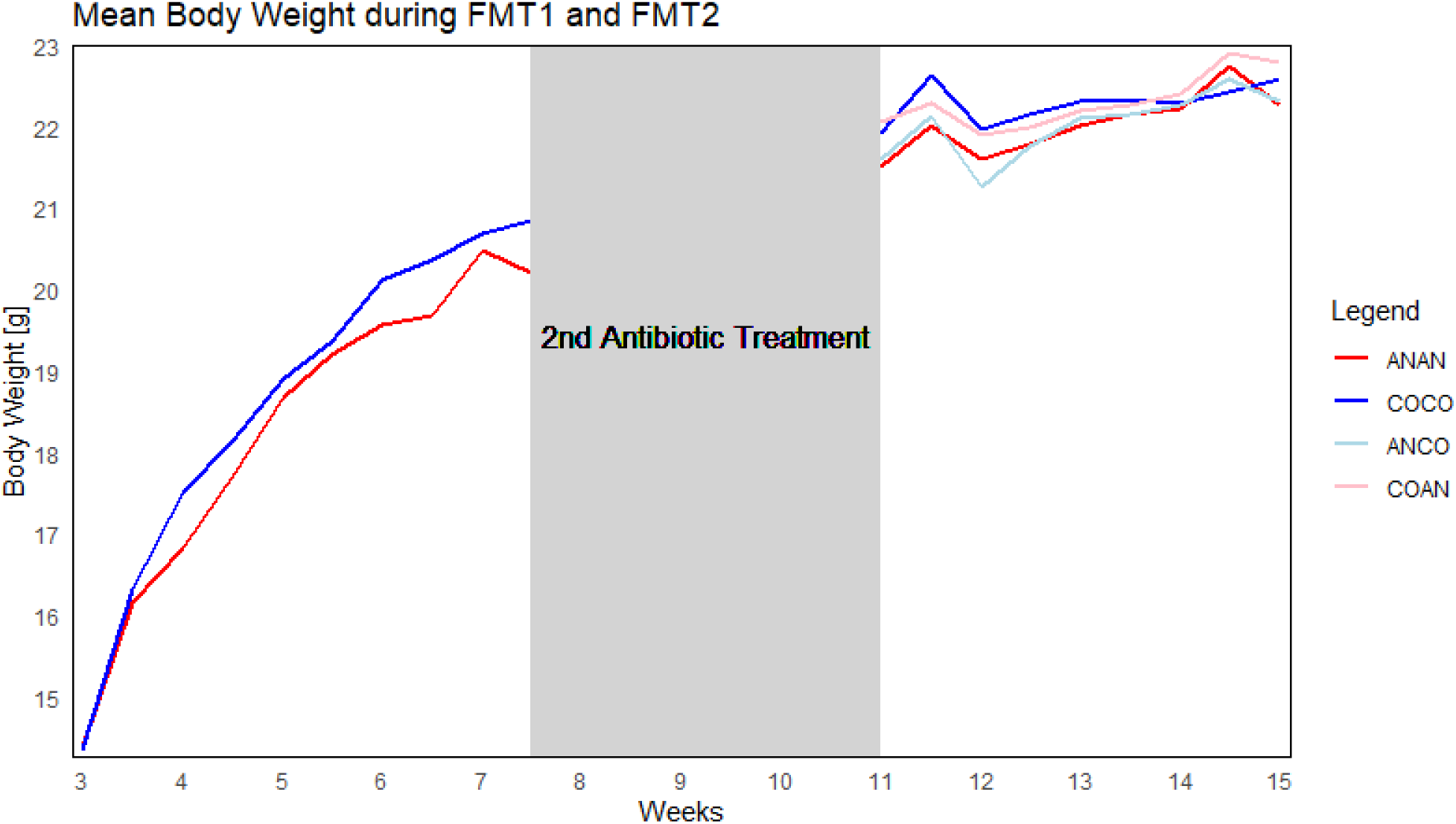
Effects of FMT from AN and CO donors on body weight development. **Left)** FMT1 (AN mice, n = 24, CO mice, n =24, (6+6 cages) for five weeks. **Middle)** three weeks of antibiotics (body weight not shown). **Right)** FMT2 shows the body weight of mice based on combined donor for FMT1 and FMT2. All mice were weighed twice a week and AUCs were calculated and evaluated by 2-way ANOVA with no statistical differences based on donor type.

*Food Intake:* Food was weighed per cage twice a week. Overall, the AN cages (after FMT1) and ANAN cages (after FMT2) had lower food intake between 1.9% and 6.9% accumulated in any comparison with other donor combinations. (For all comparisons see Supplementary Table 1). As shown in Figure 2, the food intake during FMT1 was significantly (\**p = 0.020*) lower in AN cages (n=6) compared to CO cages (n=6). There were no statistical differences between our two consecutive experiments, ExpA and ExpB, after FMT1 and hence analyzed as one.

**Figure 2.**
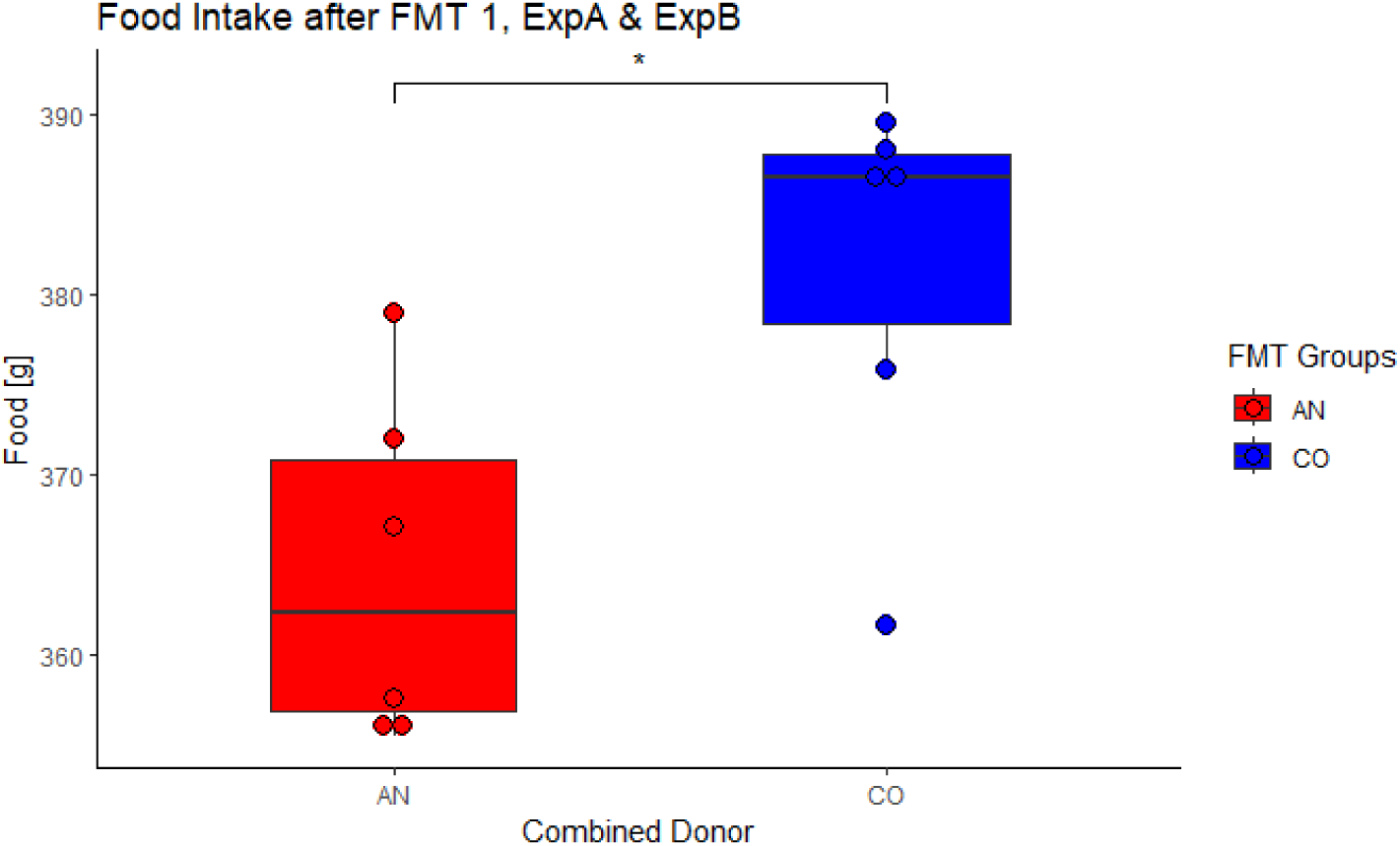
Effect of FMT1 on accumulated food intake over five weeks per cage (4 mice/cage). Statistical differences between AN cages (n=6, (mice n = 24)) and CO cages (n=6, (mice n = 24)) were evaluated by 2-way ANOVA (AN/CO: \**p* < 0.020).

However, after FMT2, ExpA and ExpB were factorially different (\**p = 0.013*) and hence analyzed separately (Figure 3A and B). After FMT2, the ANAN cages have a reduced accumulated food intake than COCO or the cross-over groups COAN and ANCO. Both cross-over groups show intermediate food intake levels. Although there is a trend, the results from the separate ExpA or ExpB did not reach statistical significance.

**Figure 3.**
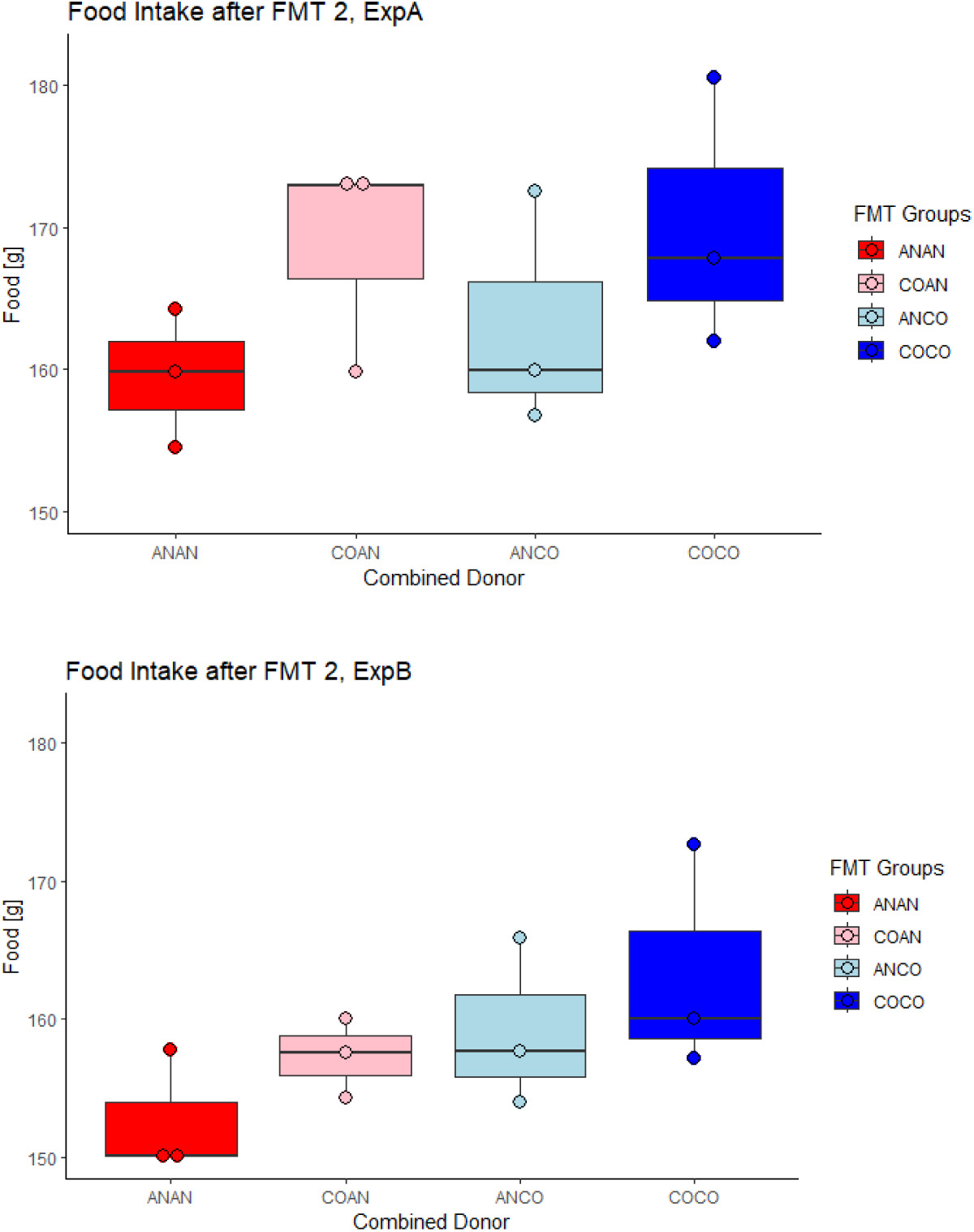
Effect after FMT2 on accumulated food intake per cage, over four weeks in a split cross-over design. Combined donor explanation: ANAN= same AN donor at FMT1 and FMT2, COAN= CO donor at FMT1 and AN donor at FMT2, ANCO= AN donor at FMT1 and CO donor at FMT2, COCO= same CO donor at FMT1 and FMT2. Statistical differences between combined donor groups and ExpA vs ExpB were evaluated by 2-way ANOVA. Since there was a significant difference between ExpA and ExpB (\**p* = 0.013), the two experiments were analyzed separately. Accumulated food intakes were then calculated and evaluated by 1-way ANOVA. A) shows ExpA (Cages n=12, (mice n = 24)) (*p* = ns). B) shows ExpB (Cages n=12, (mice n = 24)) (*p* = ns).

### Satiety-Inducing hormones are increased in AN Mice

At the end of the FMT2, the animals were killed, and serum concentrations of six satiety and appetite related hormones were measured. As can be seen from Figure 4, ANAN mice had significantly higher concentrations of PYY (increases satiety, \**p = 0.012*) and leptin (long term suppresser of food intake, *\*p = 0.019*), compared to COCO mice. Insulin (inhibits feeding) and glucagon (suppresses appetite) were on average elevated in AN mice but did not reach statistical significance. Additionally, CO mice had on average higher serum concentrations ghrelin (appetite stimulating) but did not reach statistical significance. There was no difference in active GLP-1 (promotes satiety) serum levels between groups.

**Figure 4.**
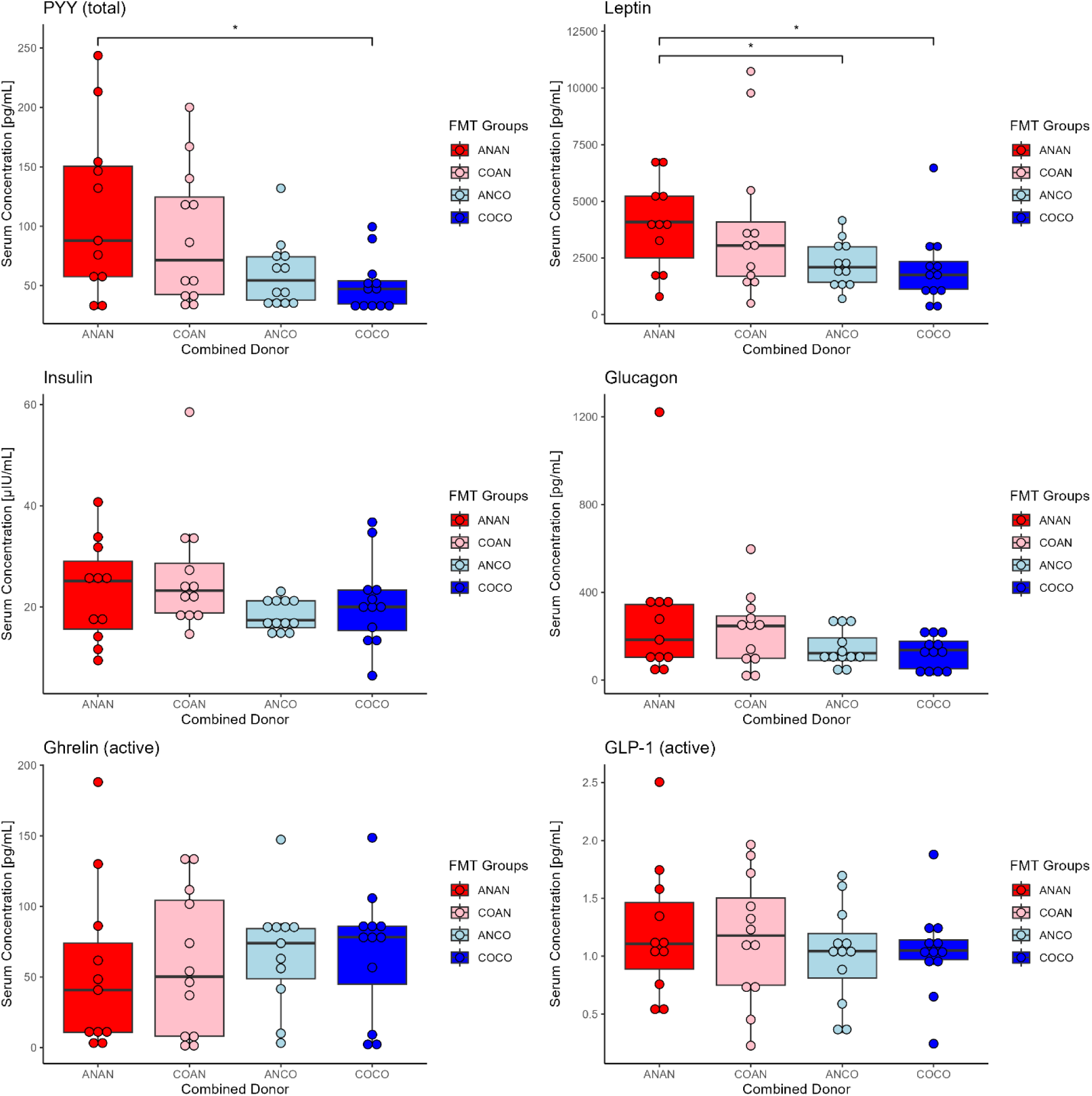
Effects of FMTs (Combined donor) on serum concentrations (pg/ml or μIU/mL) on appetite related molecules in serum. 1) PYY, 2) leptin, 3) insulin, 4) glucagon, 5) ghrelin and 6) GLP-1 (active), were measured and evaluated by 2-way ANOVA. There were significant differences in PYY and leptin serum concentrations (\**p = 0.012*, \**p = 0.012* respectively) and no significant differences in insulin, glucagon, ghrelin and GLP-1 (active).

### Humanization of Gut Microbiota in Mice after Antibiotics Treatment

We manipulated the gut microbiota (GM) of the mice with AB and humanized them during FMT1 and FMT2 (Figure 5, *p* < **0.01). Based on the similarity coefficient Jaccard index which relies on the presence and absence of ASVs, we succeeded in depleting the GM from the original mouse GM (FS1=red) with antibiotics, illustrated by the shift in clustering from red to blue (FS2=blue). After five weeks of FMT1 the humanized mouse GMs (FS3=orange) are shifted (from blue) towards the human donors (AN=pink and CO=black) and still separate from the original mouse GM (FS1=red). After a second round of AB (FS4, data points are not shown), split cross-over and four weeks of FMT2, the humanized mouse GM is shifted once again (FS5=green).

**Figure 5.**
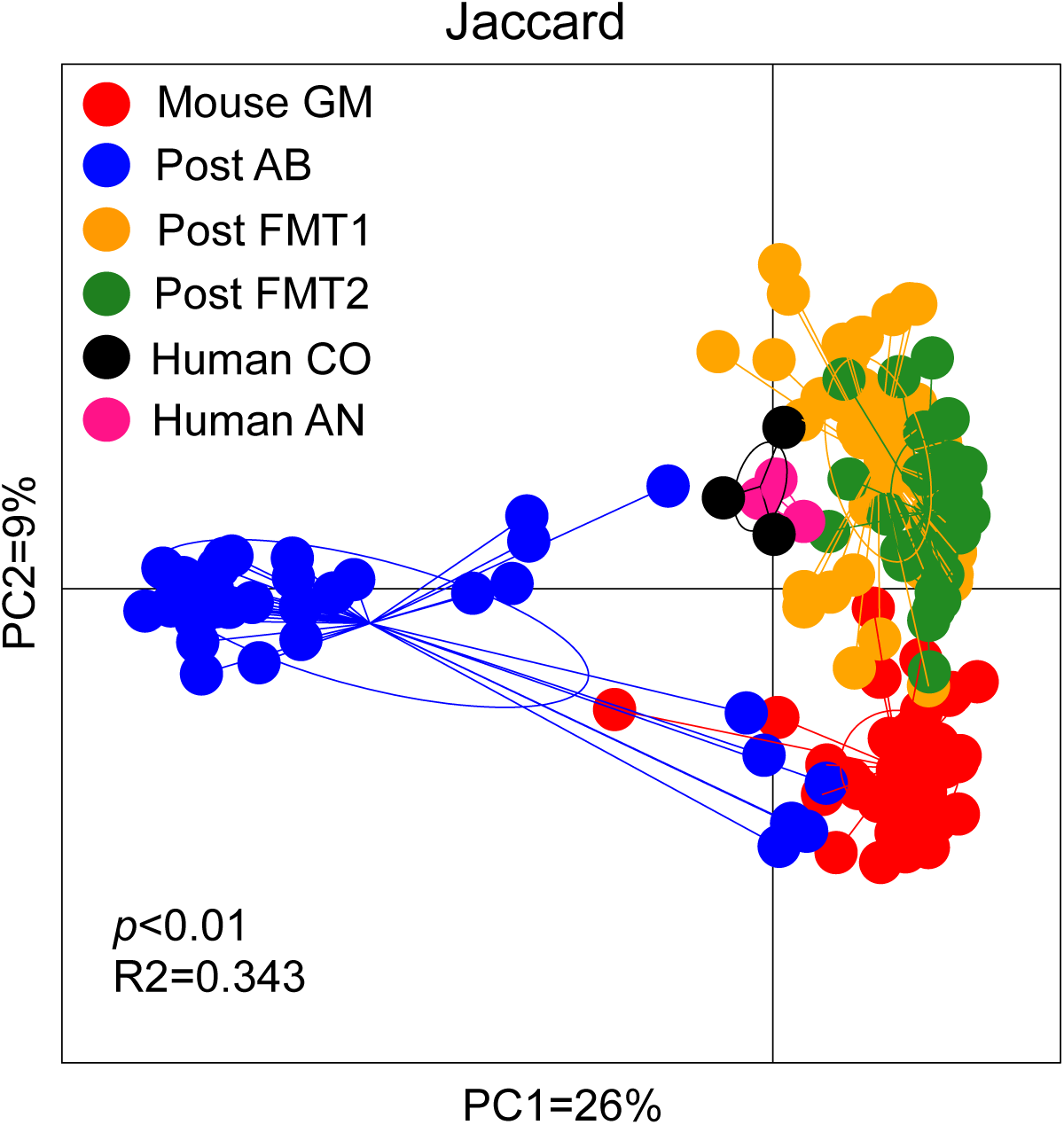
Gut microbiota of human and mice samples shown by the Jaccard index. The effect of the antibiotic treatment can be seen in the first shift in microbiota from the non-manipulated mouse microbiota (FS1, red) to the mouse GM which was taken after AB treatment (blue, FS2). After five weeks of FMT1 the humanized mouse GM (orange, FS3) shifted from blue towards the human donor (AN, pink and CO, black) while still separated from the original mouse GM (FS1, red). After the second round of AB (not shown, FS4) and four weeks of FMT2, the humanized mouse GM is shifted again (green, FS5).

After FMT1, our two consecutive experimental runs ExpA and ExpB were factorial different (\*\**p* < 0.01, Supplementary Figure 1) and thus analyzed separately. In both ExpA and ExpB the GM of the AN group was significantly different from the CO group (\**p* < 0.02, \*\**p* < 0.01, Figure 6). After FMT2, the two experimental runs were still significantly different (\**p* < 0.02, Supplementary Figure 2). However, after FMT2, there is no longer a significant difference between the now four subgroups in either experimental run (Supplementary Figure 3). There is also no statistical difference when comparing either the four subgroups regarding their last donor or when comparing only the ANAN and COCO groups (data not shown).

**Figure 6.**
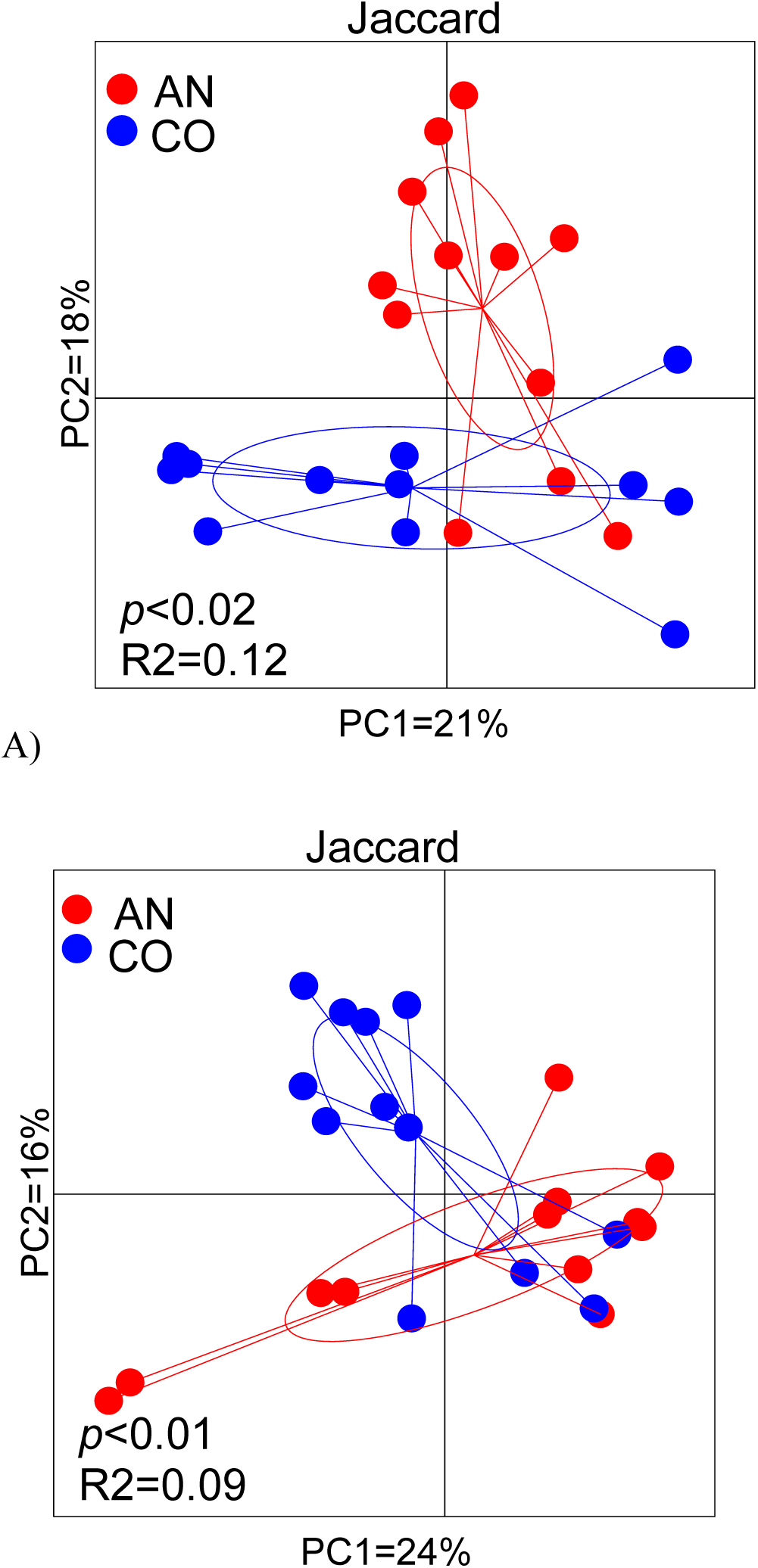
Effects of FMT1 on the mice GM in ExpA (A) and ExpB (B). The microbiota is compared by the Jaccard index. After FMT1 the mice receiving AN microbiota (red) were significantly different from CO receiving mice (blue) in both experimental runs (ExpA: \**p* < 0.02, ExpB: \*\**p* < 0.01).

We also sequenced the GM from the six human donors and the relative abundances can be seen in Figure 7 and Supplementary Figure 5. With six different donors attributed to two categories (AN or CO) there is no meaningful statistical comparison to their differences, but noteworthy GM families are *Ruminococcaceae* and *Lachnospiraceae*, as well as the genera *Faecalibacterium*, *Bifidobacterium* and *Ruminococcus*.

**Figure 7.**
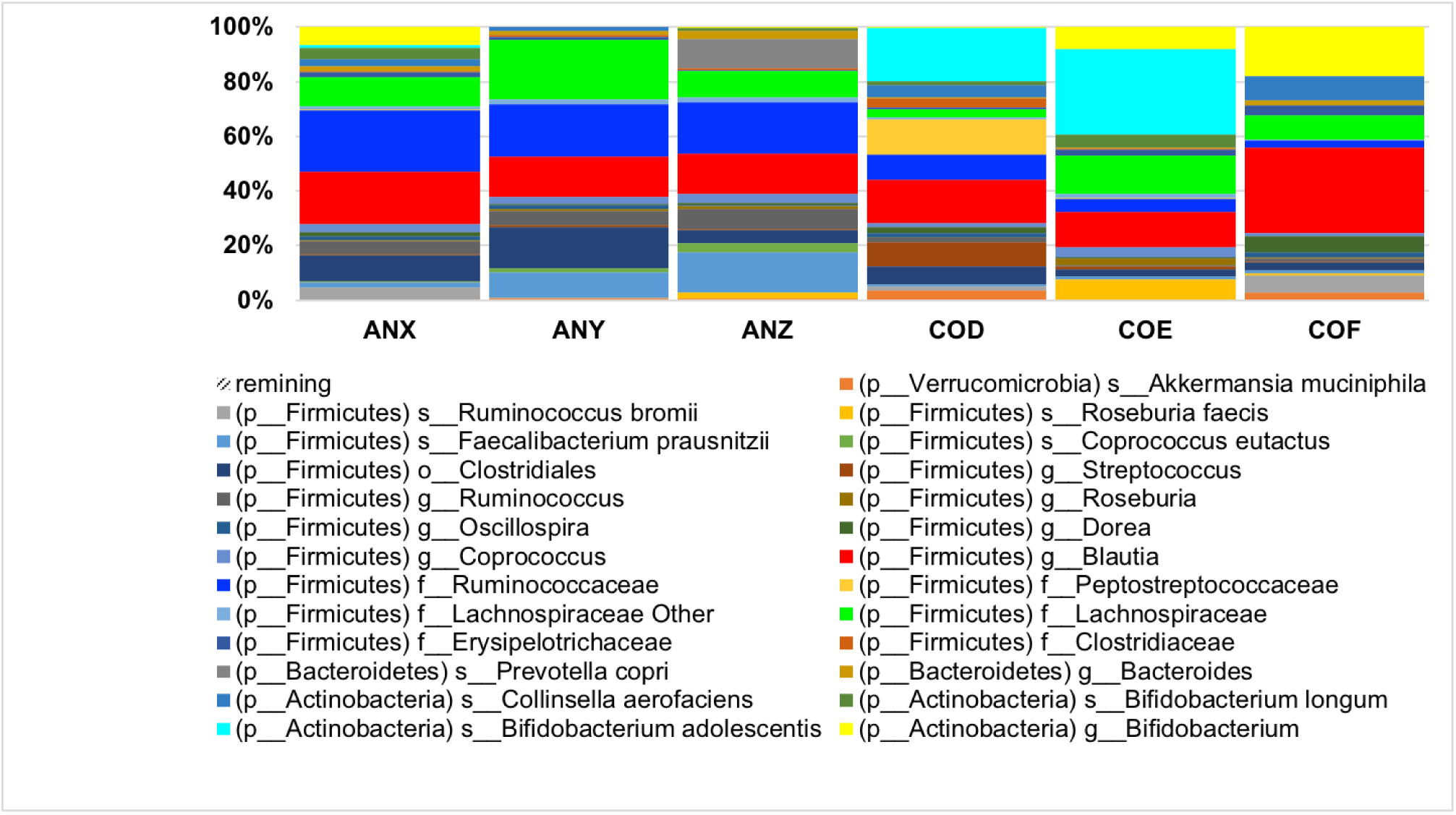
Composition of the GM of the six human donors. ANX, ANZ and ANY are the three AN donors and COD, COE and COF are the control donors. The bacterial compositions are presented in a relative correlation to each other. In the AN donors the abundances of *Faecalibacterium* (light blue), *Rumicnococcaceae* (blue) and *Lachnospiraceae* (green) are higher compared to CO donors. In two of the CO donors the abundance of *Bifidobacterium* (yellow and bright blue) is higher.

When analyzing the GM of all the mice after five weeks of FMT1, the ExpA and ExpB were factorial different (\**p* = 0.007) (Supplementary Figure 1), and the two experiments were analyzed separately. Looking into the bacterial difference in the GM, there is a significant difference in *Clostridium* abundance (Figure 8).

**Figure 8.**
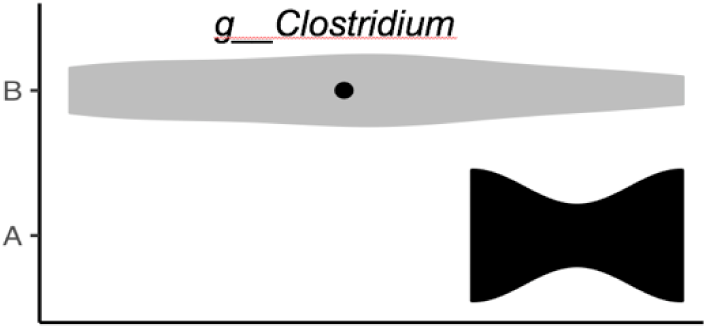
After FMT1 the abundance of *Clostridium* is significantly different between the two experimental runs ExpA and ExpB.

After FMT1, in ExpA the GM of the AN mice were significantly different from that of CO mice (Jaccard, *\*\*\*p* < 0.001) (Figure 6A). In ExpA, AN mice had a higher abundance in *Bilophila* (Figure 9). In ExpB (After FMT1), the GM of the AN mice were also significantly different from that of CO mice (\**p* = 0.004) (Figure 6B). However, there were no statistical significances when analyzed at lower taxonomic levels.

**Figure 9.**
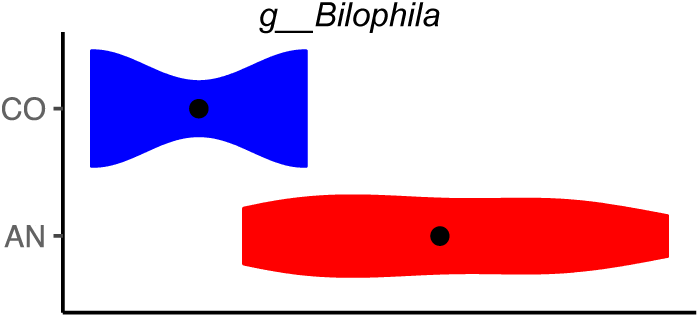
After FMT1 the microbiota of AN and CO receiving mice were compared. There is a significant difference in the abundance of *Bilophila*, which is higher in AN mice.

When analyzing the GM of all the mice after four weeks of FMT2, there were not any significant differences when analyzed either based on last donor type (data not shown) or the combined donor after FMT2 (Supplementary Figure 3). There were, however, significant differences between the two consecutive experiments ExpA and ExpB in the abundance level of *Adlercreutzia, S24-7*, *Lachnospiraceae, Ruminococcaceae, Ruminococcus, Rikenellaceae* and *Streptophyta* (Figure 10).

**Figure 10.**
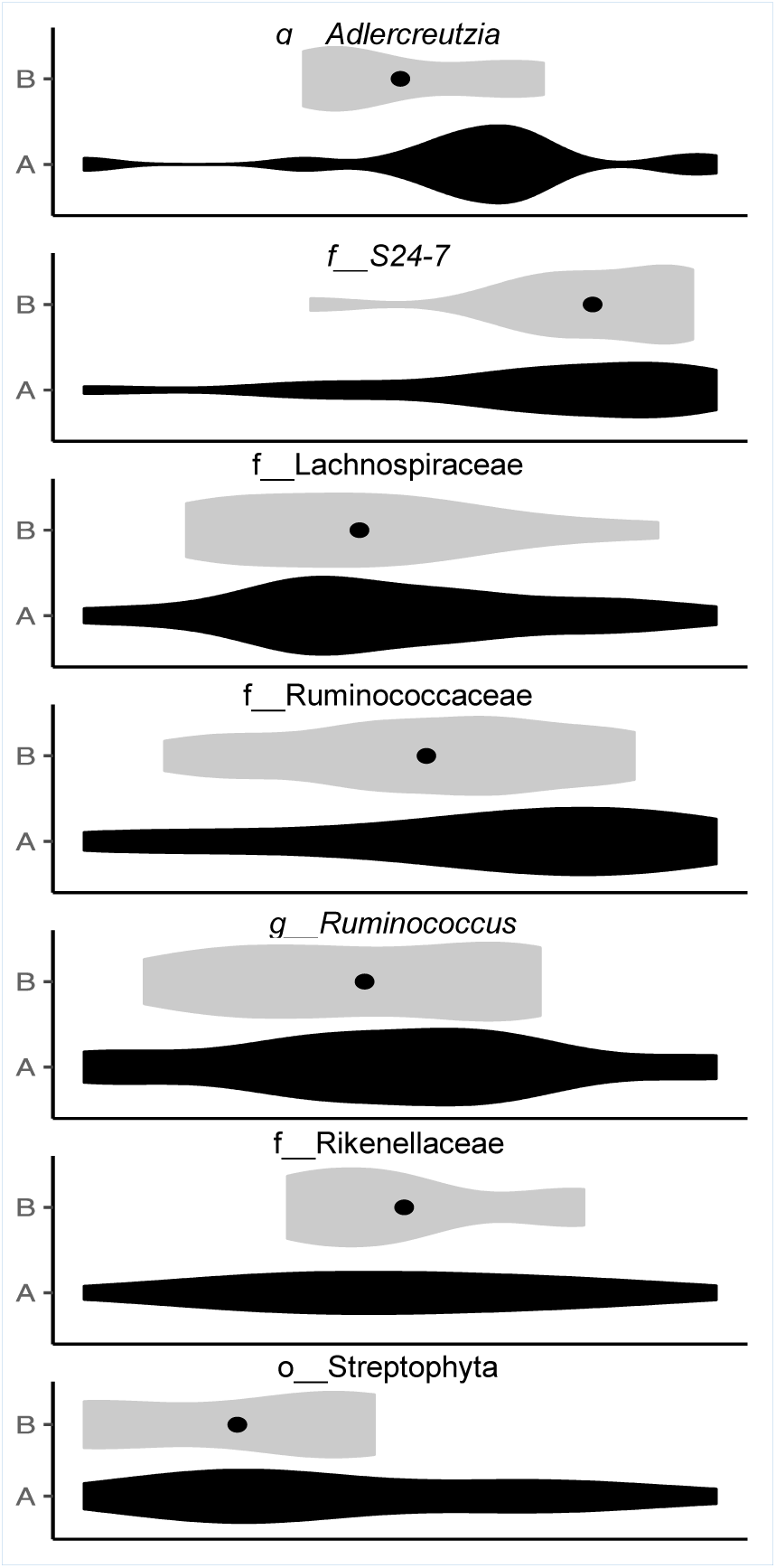
Statistically significant bacteria when comparing ExpA and ExpB after FMT2. The genera *Adlercreutzia*, *Ruminococcus*, and the families *Lachnospiraceae*, *Ruminococcaeae*, *Rikenellaceae* and the order *Streptophyta* have a higher abundance in ExpA. While *S24-7* has a higher abundance in ExpB.

### AN Mice show less active behavior during Open Field Tests

To evaluate any effect of FMTs on general locomotor activity level, the mice were subjected to open field (OF) tests. Baseline measurements were performed after the initial two weeks of AB treatment but before the first FMT. Baseline data did not show any differences in general locomotor activity between groups (data not shown).

After five weeks of FMT1, the open field data shows a factorially difference between our two consecutive experiments, ExpA and ExpB (\*\*\**p* < 0.001) and hence we analyzed them as separately (Figure 11). In ExpA, there was no differences in neither the general locomotor activity level (total distance moved) nor the total frequency of crossing the field’s subdivisions (Figure 11A). In ExpB, both the total distance (\**p* = 0.024) and frequency (\**p* = 0.024) were significantly lower in AN mice compared to CO mice (Figure 11B). After cross-over and FMT2, there were no differences in general locomotor activity and frequency in either ExpA or ExpB, when analyzing by last donor type or combined donor (data not shown).

**Figure 11.**
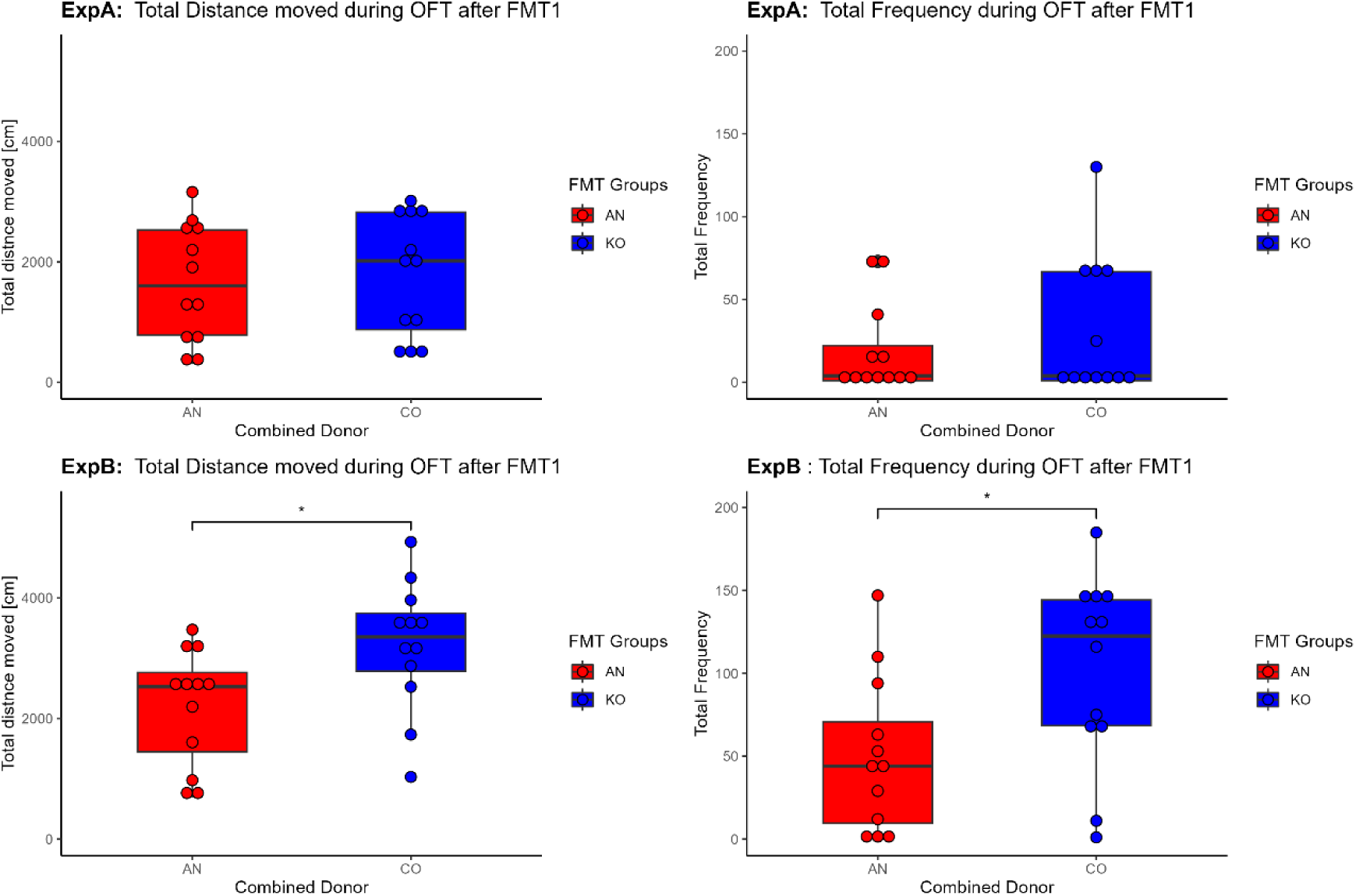
General locomotor activity. Effect on total distance moved (cm) in the open field and total frequency of crossing between the three open field subdivisions (outer, middle and center subdivision) from FMT1 compared between AN versus CO donor. In ExpA there were no differences observed. **A1)** Total distance moved in ExpA, **A2)** Total frequency in ExpA (AN, n =12) (CO, n = 12). In ExpB, we observed a significant decrease in the AN mice compared to CO mice in both total distance moves and crossing frequency. **B1)** Total distance moved in ExpB, **B2) T**otal frequency in ExpB (AN, n = 12) (CO, n = 12) (\**p* < 0.010).

Over time and regardless of FMT donors, we did observe a significant decrease in general locomotor activity (\*\*\**p* < 0.001) and total frequency (\**p* = 0.012) for both ExpA and ExpB (between baseline and FMT1). We also observed at significant decrease general locomotor activity (\*\*\**p* < 0.001) and total frequency (\*\*\**p* < 0.001) in the time between FMT1 and FMT2 (Supplementary Figure 4).

## Discussion

We have investigated if phenotypes related to Anorexia Nervosa (AN) such as lower body weight and appetite as well as altered behavior could be transferred by fecal microbiome transplants (FMT) from AN patients to antibiotic (AB) gut microbiota (GM) depleted mice. Using a split-group cross-over design, we hypothesized that we might ameliorate body weight gain, food intake, hormone levels and behavior introduced by FMT of the AN gut microbiota (AN-GM), by a second later FMT round with GM from a healthy control donor. This was done as a way of early exploring FMT as a possible therapeutic approach to rectifying AN-GM dysbiosis in humans.

We successfully depleted the indigenous mouse gut microbiotas with AB and humanized the model by grafting the human donor GM to the SPF mice as before (23). While others have used fresh donor material (19) and germ-free mice (GF) (19, 20), our approach has a few advantages. Since we used frozen donor samples with added glycerol for GM stability (24), we had easier access to enough comparable FMT material to perform repeated experiments. Furthermore, the added glycerol has the advantage of removing the need for repeated gastric gavages of the mice as they voluntarily ingest the inoculum, when it is presented. The advantage in using AB depleted SPF mice instead of GF mice depends on the questions we are trying to answer, and on the human situation we are trying to mimic (22). Fan et al. (2023) used FMTs on GF mice that received a calorie restricted diet for three weeks after two gavages of FMT two days apart, but did not report on food intake (11). They investigated weight gain, GM differences by amplicon sequencing and performed hypothalamic gene expression analysis. Hata et al. (2019) inoculated pregnant GF mice and investigated the AN characteristics in the offspring. In their experimental setup Hata et al. (2019) investigated the influence of AN-GM as a cross-generational effect, and studied mice that have developed their GM, physiology, metabolism and immunology influenced by the human GM before birth and during their early life development (19). In the study of Glenny et al. (2021), AN-GM was transferred to adolescent GF mice that had developed their physiology, metabolism and immunology in their early life without being contested by human GM. FMTs in this GF model mimics an induced change in GM, but on a naïve immunological and physiological background and the transplant in itself initiates a series of novel immune interactions (25). In our AN-GM study, we used AB depleted SPF mice, as we wanted to investigate a shift in GM on an adolescent, immune competent background. This approach mimics a possible human situation, where the GM has been changed and shifted away, from a hypothesized previously healthy state to a dysbiotic state due to changed behaviors of the patient.

After our FMT1, there was a significant decrease in accumulated food intake in AN inoculated cages compared to CO cages (Age six to eleven weeks old) ranging from 1.9% to 6.9 % less food. This corresponds well to the results reported in Hata et al. (2019) who also used BALB/c female mice, showing approximately an average of 4% less food intake during the ages four to ten weeks old. This is in contrast to Glenny et al. (2021) who reports that they observe no changes to average daily food intake between experimental groups based on the C57BL/6 mice but including both sexes of ages nine to thirteen weeks old. The discrepancies in food intake results between our and the Hata et al. (2019) study, compared to the Glenny et al. (2021) results, could be due to different mouse strain, any sex differences from the male mice that would mask changes in the females and finally the age range during observations. We, therefore, speculate that AN-GM effect on food intake might be interesting to compare directly between the sexes alone. The reduced food intake in our study does not lead to lower body weights, which corresponds to Glenny et al. (2021). Whereas Fan et al. (2023), under calorie restrictions, and Hata et al. (2019), under *ad libitum* food conditions, did show a decreased weight gain in offspring of AN inoculated mice calculated as change in body weight from baseline (11, 19, 20). The explanations for the fact that a reduced food intake does not directly lead to weight loss can be several. One explanation could be that the mice, in our study, had a lower energy expenditure, e.g. by simply moving less (26), and the differences only becomes clear if using calorie restriction as in Fan et al. (2023) (11). Another explanation could be a changed energy extraction efficiency facilitated through the GM or transit time through the GI as the energy that is extracted from the food has been shown to depend on GM composition and transit time (27, 28). One could also speculate how calorie restriction is best implemented in an AN mouse model as low-calorie *ad libitum* food or food restriction by periodically removing food entirely from the cage mimicking uneven AN eating patterns. In Fan et al. (2023) they calculated food intake of a 3 day baseline before FMTs and reduced the total amount of food available to 30% less than that (11).

After our second round of AB treatment, split cross-over and FMT2, there is a trend of decreased food intake in the pure ANAN cages compared to pure COCO, that have only received GM from one single donor through both FMT1 and FMT2. In the cross-over groups, (ANCO or COAN) with different donors in FMT1 and FMT2, intermediate food intake levels were shown, mostly influenced by the last donor type. Considering food intake alone, we can influence CO mice (FMT1) to eat less by inoculating with an AN-GM in FMT2. That also seems to be the case in the opposite situation, in which we influence AN mice (FMT1) to eat more by inoculating them with a CO-GM in FMT2. This suggests some rescue and amelioration of the AN characteristic, food intake, by FMT from a healthy donor.

In order to investigate if this correlation between FMT donors and food intake could be explained by changed appetite and satiety signals in the blood, we measured six relevant biomarkers in the serum of the mice at termination after FMT2. Although, it would have been very informative to know the level of these biomarkers already after FMT1, the amount of serum needed for the assays prevented us from measuring any before termination. This should be considered in future experimental setups. Hata et al. (2019) suggests that endocrine biomarkers such as PYY (peptide tyrosine tyrosine), ghrelin and leptin are to be considered the signature of chronic food restriction (29). Our results do show significantly higher serum concentrations of PYY and leptin in ANAN mice compared to COCO after FMT2. PYY functions as an appetite suppressing gut hormone and circulating PYY increases satiety, inhibits gastrointestinal motility, pancreatic hormone secretion, and decreases food intake (30). Likewise leptin is a mediator of long-term regulation of energy balance, suppressing food intake (31). Insulin and glucagon were on average elevated in AN mice but did not reach statistical significance. Glucagon reduces body weight and adiposity by suppressing the appetite hormone that increases during fasting. Glucagon also acts in the liver to reduce meal size, the signal being relayed to the brain via the Vagus nerve (32). It also stimulates the synthesis and secretion of ghrelin. Again, in the cross-over groups (ANCO or COAN) with different donors in FMT1 and FMT2, we show intermediate biomarker levels that are mostly influenced by the last donor type. We did not reach statistically significance on ghrelin or GLP-1, however the mean levels of ghrelin, that increases appetite (29), are higher in both CO groups (ANCO and COCO) compared to both AN groups (ANAN and COAN). The elevated serum PYY levels in our AN mice corresponds to reports of PYY and leptin contributing to lower food intake and a higher feeling of satiety in AN patients (33). PYY acts on the brain by inhibiting the orexigenic neuropeptide Y (NPY) (34). Inhibiting NPY increases the activation of neurons expressing the anorexigenic neuropeptide pro-opiomelanocortin (35). This suggests that the activity of the AN-GM transferred to our mice affects the gut-microbiota-brain axis by inducing an increased PYY signal.

In our study we find that leptin levels of AN mice are increased compared to CO mice, which suggests that AN mice felt increased satiety. However, it has been reported that in fasting AN patients the serum leptin concentrations are significantly lower than normal (36–38). The reason for this discrepancy is unclear, however, leptin is mainly secreted from subcutaneous fat (39) which is considerably reduced in adult AN patients (40), resulting in reduced leptin levels in AN patients. Even though the body weight of our mice were the same for AN mice and CO mice, the amount adipose tissue might be different. In Fan et al. (2023), FMT from human AN donors resulted in increased abundance of Ucp1, Elovl3 and Pgc1α mRNA in inguinal fat of AN-transplanted mice, indicating enhanced adipose tissue thermogenesis in this group of mice. The time frame of which we observe the mice and the influence of the change in GM to AN-GM must also be considered a short amount of time compared to the human AN situation. The serum insulin and glucagon levels were also increased in the AN mice, however we did not fast the mice before blood sampling and is important for future studies as insulin is rapidly increases in response to glucose levels (41). There were no significant differences in serum GLP-1 or ghrelin, but ghrelin appears slightly decreased for AN mice compared to CO mice as would be expected as leptin is known to inhibit secretion of ghrelin (36). The GLP-1 gut hormone is co-secreted with PYY from intestinal L-cells following a food intake to mediate post-prandial satiety and has been shown in human studies to have a synergistic effect on food intake (42–44).

To comprehensively examine the impact of the FMTs on the microbiome of the humanized mice, we sequenced fecal samples collected at different times. We analyzed the microbiota using the Jaccard index, which considers the presence and absence of ASVs, and subsequently, we assessed statistical differences in bacterial genera at lowest possible taxonomical level.

A first observation was the successful GM shift of the naïve mouse GM through the antibiotic treatment. Following FMT1 for five weeks, the humanized mouse GM underwent a shift towards that of the human donors, while remaining distinct from the original mouse GM. Following the second round of AB treatment and four weeks of FMT2, the humanized GM underwent another shift.

After FMT1, the two experimental runs, ExpA and ExpB, were significantly different and were therefore analyzed separately. In both experimental runs, we observed significant differences between AN and CO mice based on the Jaccard Index. However, after FMT2, we did not observe any statistically significant differences in GM when comparing the combined donor types (ANAN, COCO, ANCO, COAN). Additionally, no significant differences were found when comparing groups based on their last donor types (ANAN and COAN vs. COCO and ANCO) or when comparing groups that received only one donor type during both FMTs (ANAN vs. COCO). This is possibly due to the lower group sizes after cross-over. Also, FMT2 is one week shorter than FMT1 and given an expected GM establishment period the FMT2 might have to be extended in the future.

We also investigated the bacterial composition of the human donor GM and the humanized mice GM and focused on genera that showed statistical significance. Among the six human donors, we noted differences in the abundance of *Ruminococcaceae*, *Lachnospiraceae* and *Faecalibacterium*, which were higher in AN donors compared to CO donors. Conversely, CO donors had a higher abundance of *Bifidobacterium*. *Lachnospiraceae* have been associated with weight gain following AB treatment and increased blood glucose levels, possibly indicating a link to energy utilization and rapid weight gain (45–47). Higher *Lachnospiraceae* abundance at hospital admission has also been shown to be an indicator to predict shorter inpatient treatment duration in a human anorexic study (15). In Fan et al. (2023), characterization of GM of 77 AN patients did show lower abundance of *Lachnospiraceae.* Similarly, previous research has found that *Faecalibacterium* abundance, at hospital admission, is lower in AN patients, and its level increases significantly with weight gain and improvement of AN (15). Fan et al. (2023) also reported *Faecalibacterium* to have lower abundance in AN patients compared to healthy controls. *Ruminococcaceae* has also been described to be altered in human AN patients and influenced by weight gain (15).

The humanized mice GM after FMT1 in ExpA and ExpB showed significant differences, with *Clostridium* being notably different in abundance between the two experimental runs. *Clostridium* species have been implicated in the regulation of eating behavior and neuropsychiatric symptoms (48), which could explain the variability observed in ExpA and ExpB.

Moreover, when comparing the GM of AN and CO mice, we found higher levels of *Bilophila* in AN mice of ExpA after FMT1. In a weight-loss surgery study, *Bilophila* has been associated with weight loss and a reduction in body fat (49).

Following FMT2, there were no statistically significant bacterial differences when analyzing between combined donor types. However, our investigation revealed significant differences between ExpA and ExpB in the abundance levels of *Adlercreutzia, S24-7, Lachnospiraceae, Ruminococcaceae, Ruminococcus, Rikenellaceae* and *Streptophyta*. Considering that *Lachnospiraceae*, *Ruminococcaceae* and *Ruminococcus* have been previously shown to be altered in human AN patients and influenced by body weight (15, 50), these varying concentrations of bacteria could contribute to the differences between the two experimental runs.

To evaluate any effect of FMTs on general locomotor activity (GMA) level, the mice were subjected to open field (OF) tests. We did observe a small statistically significant decrease in AN-mouse activity and frequency of crossing the subdivisions of the OF after FMT1 in ExpB (ten to eleven weeks old). However, this was not observed in ExpA. Hata et al. (2019) showed no significant differences in GMA (20 min), but their AN-GM mice also spent significantly more time in the peripheral subsquares than their control mice of the same age (\*\*\**p =* 0.0009) (19). In our study, any behavioral changes due to FMT leading to decreased activity level and energy expenditure might explain the maintained body weight in the AN mice despite their decrease in food intake. This is in contrast to anorexia in humans, which often is correlated to increased activity excess exercises (51). It would therefore be interesting to measure GMA continuously in the home cages or possibly combine FMT with exciting AN mouse model of exercise (52) to observe any accelerated weight loss. After FMT2, there were no differences in GMA or frequency based on FMT donors. We did however, by serendipity, observe a clear effect of time and/or our repeated measurements in the OF, leading to decreased GMA and frequency in crossing the subdivisions, that might mask more subtle GM-induced behavioral changes. We cannot discern whether the decreases were due to FMT procedures, repeated exposures resulting in decrease of novelty, our antibiotic treatment, or age related as our mice get older. We observed a significant decrease in GMA (\*\*\**p* < 0.001) and total frequency (\**p* = 0.0124) for both ExpA and ExpB when comparing start (baseline) and end of the experiment, which shows that time and repeated measurements are important factors that must be considered when performing behavioral experiments. To overcome repeated exposures to the same environment, it would be interesting to use a non-interfering system to measure spontaneous physical activity such as the system developed by Poffé et al. (2018) (53).

In conclusion, our study suggests that gut microbiota transplants derived from AN patients may contribute to the pathological characteristics of the AN, by altering food intake, through the gut-brain axis, in immune competent AB treated mice. The decreased food intake and appetite biomarkers can be ameliorated by a second FMT with gut microbiota from a healthy donor. These results may ultimately contribute to the development of supportive treatment with FMT in anorexia nervosa.

## Methods

### Subject Recruitment and Sample Collection

For this study we enrolled three female patients diagnosed with restricting-type anorexia nervosa (AN), admitted to Mental Health Centre Ballerup, and additionally, three healthy female controls (CO) with no history of any eating disorders or digestive diseases such as IBD or IBS (Table 1). None of the donors had used antibiotics three months prior to study participation. The study protocol was approved by the local ethics board (id: H-15012537; addendum 77106) under the PROspective Longitudinal all-comer Eating Disorder study (PROLED) study. Written informed consent was obtained from all participants before enrolment in the study.

**Table 1.** Characteristics of anorexia nervosa patients and healthy controls for feces collection.

| **Donor** | **Age** | **BMI, kg/m<sup>2</sup>** |
| --- | --- | --- |
| **ANX** | 18 | 16.90 |
| **ANY** | 21 | 15.42 |
| **ANZ** | 22 | 16.89 |
| **Mean ± SE** | **20.33 ± 1.20** | **16.43 ± 0.49** |
| **COD** | 25 | 19.70 |
| **COE** | 30 | 23.60 |
| **COF** | 27 | 19.83 |
| **Mean ± SE** | **27.33 ± 1.45** | **21.04 ± 1.28** |

A fresh fecal sample was collected from the three (AN) donors (ANX, ANY, and ANZ) and from the three control (CO) donors (COD, COE, and COF). Body mass index (BMI) and age of the six participants are shown in Table 1. At the time of sample collection, the fresh stools were divided into four tubes and subsequently stored at -80 °C until further processed for animal experiments.

### Animals and Housing

48 specific-pathogen free (SPF) female BALB/cJTac mice (Taconic Biosciences, Ejby, Denmark), aged three to four weeks at arrival, were randomly distributed to cages and housed at the Association for Assessment and Accreditation of Laboratory Animal Care (AAALAC)-accredited barrier protected facility of the Faculty of Health and Medical Sciences, University of Copenhagen, Frederiksberg, Denmark. The sample size was founded on a power analysis, which revealed a sample size of ten per donor group with 90 % power and 5 % significance level based on comparable results from Hata et al. (2019) (19).

The facility maintained a temperature of 22 °C (+/- 2 °C), a humidity of 55% (+/- 10%), 12/12-hour light/dark cycle (lights on from 7 a.m., summertime), and 15-20 air changes per hour. Upon arrival, all mice were randomized blindly, earmarked and then housed four per open cage (Techniplast, 1290D Eurostandard type III, Scanbur A/S, Karlslunde, Denmark), with Tapvei® aspen bedding, nesting material Enviro-dri®, a cardboard Smart Home shelter, a fresh 1 x 1 x 5cm Tapvei® aspen chewing block, fresh cotton Nestlets, a cardboard Mini Fun Tunnel with changing of bedding, and a clear plastic tunnel (Brogaarden, Lynge, Denmark) with water and feed *ad libitum*. The bedding was only changed at the end of each AB treatment for maximum bacterial exposure. The mice received the maintenance diet Altromin 1324 feed (Brogaarden, Lynge, Denmark) consisting of 11% fat, 24% protein, and 65% carbohydrates. According to Federation of European Laboratory Animal Science Associations (FELASA) guidelines (54) routine health monitoring of the mice revealed no listed pathogens.

### Animal Ethics

This study was conducted according to the Danish Act on Animal Experimentation (BEK nr 2028 af 14/12/2020) and the EU directive “on the protection of animals used for scientific purposes” (2010/63/EU). The Animal Experimentation Inspectorate, Ministry of Food, Fisheries, and Agriculture, Denmark approved this study (license no. 2017-15-0201-01262). The study was planned according to PREPARE guidelines (55), and it will be reported according to ARRIVE guidelines (56, 57).

### Antibiotic Treatment

All mice were treated with Ampicillin (Sigma-Aldrich, St. Louis, MO, USA) in the drinking water (1 g/L) twice, first for two and later for three weeks, as previously described (58). The mice received the water with ampicillin *ad libitum* fresh every three days. The antibiotic and dose were chosen as it has previously been shown to consistently reduce gut bacterial density and facilitate transfer of donor GM (59).

### Fecal Matter Transplant (FMT)

Before FMT, the fecal samples from patients and controls were thawed on ice and subsequently suspended 1:5 in 50% glycerol. 300 μL of the suspension was divided into 2 mL Biosphere® SafeSeal tubes, one tube per FMT. The samples were stored at -80 °C until thawing in ice for administration. FMT was performed once a week on all mice, where 50 μL of suspension was applied at room temperature to the mouth of the mice with a sterile 1 mL disposable syringe.

### A “split-group cross-over” Study Design (FMT1 and FMT2)

There were two consecutive FMT phases (FMT1 and FMT2) in this study performed on the same mice (n=48), with a partial cross-over midway to discern any changes, compared to non-cross-over treatment “clean” groups. All mice were weighed twice a week and food intake per cage was calculated weekly. We performed this as two identical series (A and B) in parallel staggered one week with 24 different mice in each experiment (ExpA, n = 24 and ExpB, n = 24). For an overview see Table 2.

**Table 2:** A “split-group cross-over” study design (FMT1 and FMT2). After two weeks of antibiotic treatment cages were randomly distributed to either experiment A (ExpA) or repeat experiment B (ExpB). **ExpA:** The six cages received one of each of the six human donors (three AN or three CO) (four mice/cage, total 24). After five weeks of FMT1, all mice were again subjected to antibiotic treatment (three weeks). At the “split-group cross-over” all cages were subdivided into two new cages (all cages, two mice/cage) either receiving GM from the exact same donor as during FMT1 (ANAN and COCO) or from the cross-over donor (ANCO and COAN) at random. **ExpB:** Exactly, repeating ExpA, but a week delayed for practical reasons.

| Experiment | AB treatment | FMT1 |  |  |  | AB treatment | FMT2 / Cross-over |  |  |  |  |
| --- | --- | --- | --- | --- | --- | --- | --- | --- | --- | --- | --- |
|  |  | Donor Type | FMT | #mice | #cages |  | Donor Treatment | Comb. Donor | FMT | #cages | #mice |
| **ExpA** | 2 weeks | AN | 5x | 12 | 3 | 3 weeks | AN | ANAN | 4x | 3 | 6 |
|  |  |  |  |  |  |  | CO | ANCO | 4x | 3 | 6 |
| **ExpA** | 2 weeks | CO | 5x | 12 | 3 | 3 weeks | CO | COCO | 4x | 3 | 6 |
|  |  |  |  |  |  |  | AN | COAN | 4x | 3 | 6 |
| **ExpB** | 2 weeks | AN | 5x | 12 | 3 | 3 weeks | AN | ANAN | 4x | 3 | 6 |
|  |  |  |  |  |  |  | CO | ANCO | 4x | 3 | 6 |
| **ExpB** | 2 weeks | CO | 5x | 12 | 3 | 3 weeks | CO | COCO | 4x | 3 | 6 |
|  |  |  |  |  |  |  | AN | COAN | 4x | 3 | 6 |

After two weeks of antibiotic treatment all mice were subjected to FMT1: For five consecutive weeks mice (four mice/ cage) were given a FMT from one of six human donors, from either three AN (Anorexic) and three CO (Controls). After a second antibiotic treatment (three weeks) the mice were subjected to the second phase. In the FMT2, we moved two mice randomly per cage to new cages, effectively doubling our number of cages, with now two mice/cage. During FMT2 mice received either exactly the same human donor as during FMT1 (ANAN or COCO cages) or they received a cross-over donor (COAN or ANCO).

### Sampling and Behavioral Recording

We collected mouse fecal pellets from all mice; before and after the first AB treatment and after both FMT1 and FMT2 phases. Figure 12 summarizes all experimental procedures performed in both ExpA and ExpB, data collection and sampling in a timeline. Open field data collection was done before and after FMT1 and again after FMT2. At termination we collected serum and additional tissues.

**Figure 12.**
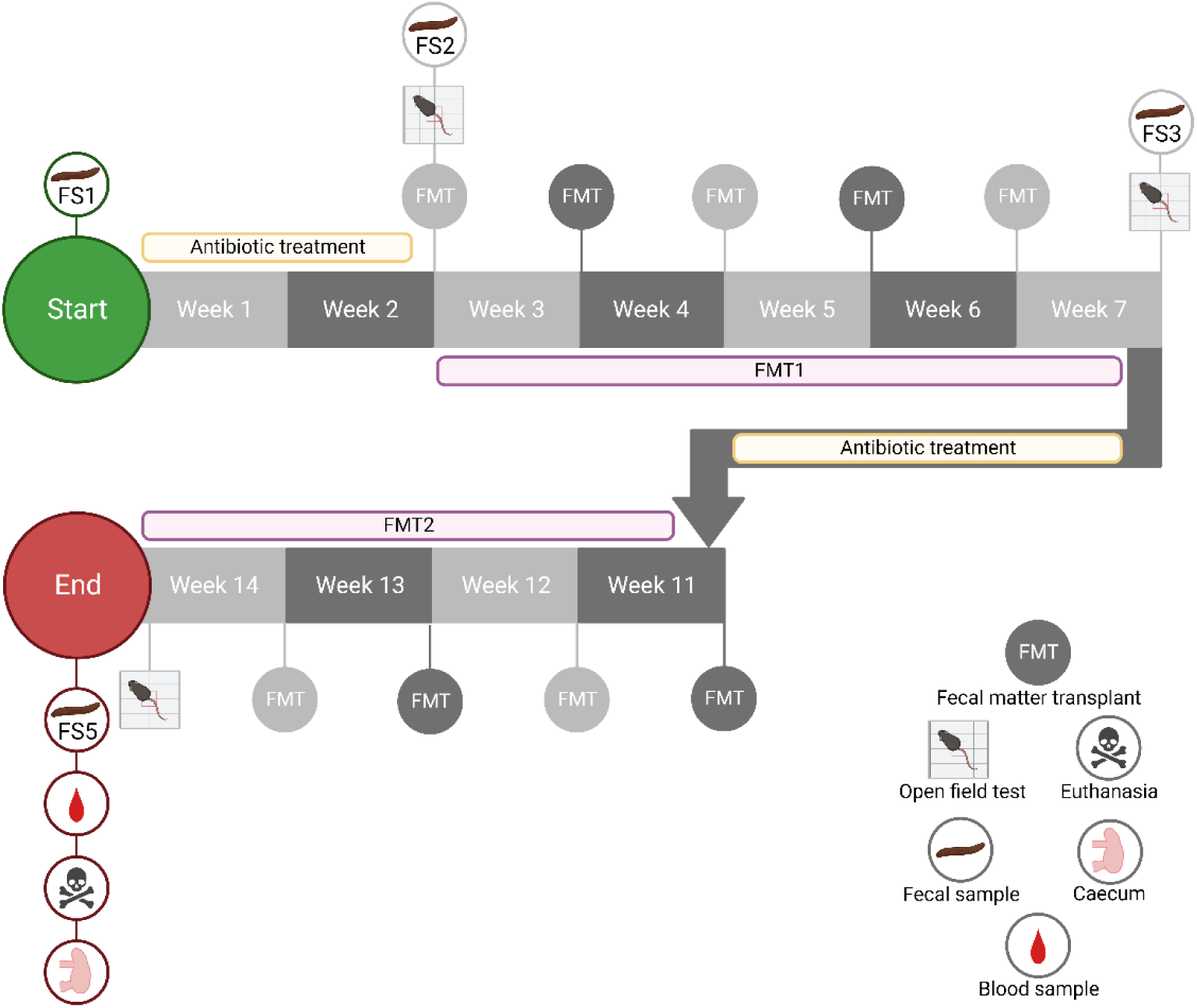
Sampling and behavioral recording in a timeline. **Start:** FMT1: Before AB treatment we collected fecal samples from all mice (FS1). All mice were subjected to two weeks of AB treatment and FS2 was collected, and the first open field behavior recorded pre-FMT1. After five weeks of FMT1, with either AN- or CO-GM, FS3 was collected and repeated open field data was recorded. After three weeks of AB treatment and split-group cross-over, mice were subjected to FMT2 for four weeks. The final open field data was collected at the end of FMT2. At the **End:** The final FS5 was taken, and mouse blood and additional tissues were collected. *Created with BioRender.com. Publication code: HM25XADTL4*

### Open Field Behavioral Test

To assess general locomotor activity level the mice were tested in an open field arena. The mice were habituated to the test room for maximum 30 minutes between 13.00 and 16.30, then in random order placed individually in the arena (48×48×28 cm). A camera placed above the arena recorded the mice activity for ten minutes. The software Ethovision XT version 13 (Noldus Information Technologies, The Netherlands) was used to sub-divide the open field arena virtually into a center zone, eight middle zones, and 16 outer zones as previously described (60) and tracking the animals’ movements in the arena. The 25 total zones are merged into three subdivisions, an outer, middle and center subdivision, on which the analyses were done. Variables used for analysis were the total distance moved (cm), mean velocity (m/s) and time and number of entries in the three different subdivisions (frequency) of the arena. Before each trial, the arena was disinfected with 70% ethanol.

### Appetite related Biomarkers in Serum

Serums were diluted 1:2 in metabolic assay working solution (MSD®, Rockville, MD, USA) and GLP- 1, glucagon, insulin, leptin, ghrelin, and PYY levels were analyzed using the U-PLEX® Mouse Metabolic Group 1 multiplex immunoassay (MSD®), following the manufacturer’s protocol.

### Gut microbiota Analysis

The DNA was extracted from fecal pellets using the Bead-Beat Micro AX Gravity kit (A&A Biotechnology, Gdynia, Poland, Cat. # 106-100-M1), following the manufacturer’s protocol. The isolated DNA was stored at -80°C until DNA library construction.

Gut microbial composition was examined by sequencing near full-length 16S rRNA gene amplicon using GridION (Oxford Nanopore Technologies, Oxford, UK) as previously described (61).

Briefly, the sequencing library was constructed using custom two step PCR for amplification of 16S rRNA gene and samples barcoding. PCR 1 reactions containing 12 μL of PCRBIO Ultra Mix (PCR Biosystems Ltd, London, UK), 6 μL of nuclease-free water, 2 μL of primer mix (5 μM), and 5 μL of genomic DNA (∼1 ng/μL) bringing the volume to a total of 25 μL were run on a SureCycler 8800 Thermal Cycler (Agilent Technologies, Santa Clara, CA, USA). The PCR program consisted of initial denaturation at 95 °C for 5 min and two cycles of denaturation 95 °C for 20 seconds, touch-down at 48 °C for 30 seconds, annealing at 65 °C for 10 seconds, and extension at 72 °C for 45 seconds, followed by final elongation at 72 °C for four minutes.

After PCR 1, the PCR products were cleaned using AMPure XP binding beads (Beckman Coulter Genomic, Indianapolis, IN, USA). The AMPure XP binding beads (Beckman Coulter Genomic, Indianapolis, IN, USA).

Following cleaning, a second PCR step was performed to barcode PCR amplicons. PCR reactions containing 12 μL of PCRBIO Ultra Mix (PCR Biosystems Ltd, London, UK), 10 μL of clean PCR 1 product, and 2 μL of barcoding primers (10 μM) bringing the volume to a total of 25 μL were run on a SureCycler 8800 Thermal Cycler (Agilent Technologies, Santa Clara, CA, USA). The PCR program was as follows: denaturation at 95°C for 2 minutes, 33 cycles of denaturation at 95°C for 20 seconds, annealing at 55°C for 20 seconds, and extension at 72°C for 40 seconds, followed by final extension at 72°C for four min.

After gel electrophoresis, 5 μL of each PCR 2 products were pooled together and subsequently cleaned with Binding Beads (AMPure XP, Indianapolis, IN, USA). The cleaned and pooled PCR products were eluted in 40 μL of nuclease-free water. The pooled and barcoded amplicons were completed for library preparation for GridION sequencing using the 1D genomic DNA by ligation protocol (SQK-LSK110) as previously described in Arildsen et al. (2021) (62). Approximate 0.2 μg of amplicons were used for the initial step of end-prep, and 40 ng of prepared amplicon library was loaded on a R9.4.1. flow cell.

### 16S Gene Amplicon Sequencing and Data Processing

The 16S rRNA gene amplicon sequencing and data processing were performed as described in Arildsen et al. (2021) (57). The data generated using GridION was collected using MinKNOW software v22.10.7 (Oxford Nanopore Technologies, Oxford, UK). The Guppy v6.2.8 basecalling toolkit was used to base call raw FAST5 to FASTQ (Oxford Nanopore Technologies, Oxford, UK). The abundance table was generated from raw FASTQ files using the Long Amplicon Consensus Analysis pipeline (LACA, GitHub: https://github.com/yanhui09/laca). Taxonomy assignment of quality-corrected reads was performed against the SILVA database (63).

The subsequent analysis was conducted using the Qiime2 (v2020.8.0) bioinformatic platform (Bolyen et al., 2019). Alpha diversity (observed species index) and beta diversity (PCoA based on Bray-Curtis and Jaccard distance matrices) were calculated through rarefaction to 3,000 reads per sample. In total, 19 samples collected fewer reads than 3,000 and were thus excluded from the analysis. Permutational multivariate analysis of variance (PERMANOVA) was used to test differences between the three groups based on Bray-Curtis and Jaccard distance matrices. Analysis of Composition of Microbiomes (ANCOM) was used to identify taxonomic groups that are differentially abundant across tested categories.

### Other Statistics

Food Intake and serum biomarkers were analyzed with one or two-way ANOVA with experimental run (ExpA vs ExpB) and FMT donor type or combined donor type as factors. For body weights, AUCs were calculated and evaluated by 2-way ANOVA. When significant we applied Tukey’s HSD. General locomotor activity and frequency evaluated by paired t-test (2-tailed).

## Acknowledgements

We would like to thank all the human donors. Denitsa Vladimirova Stefanova for excellent laboratory work on the sequencing. Data were generated through accessing research infrastructure at the University of Copenhagen, including FOODHAY (Food and Health Open Innovation Laboratory, Danish Roadmap for Research Infrastructure.)

## Disclosure of Interest

Sina Maschek

SM declares no conflict of interest.

Theresa Helena Østergaard

THØ declares no conflict of interest. Lukasz Krych

LK declares no conflict of interest.

Line Fisker Zachariassen

LFZ declares no conflict of interest. Dorte Bratbo Sørensen

DBS declares that he has collaborated with pharmaceutical industry and received funding from this source as described on https://research.ku.dk/search/result/?pure=en/persons/114880

Caroline M. Junker Mentzel

CMJM declares no conflict of interest. Axel Kornerup Hansen

AKH declares that he has collaborated with pharmaceutical industry and received funding from this source, as well as he is the owner of a diabetes related patent as described on https://ivh.ku.dk/english/employees/?pure=en/persons/107126.

Magnus Sjögren

MS declares no conflict of interest. Kenneth Klingenberg Barfod

KKB declares no conflict of interest.

## Availability of Data and Materials

## References

1. Steinhausen H-C, Jensen CM. Time trends in lifetime incidence rates of first-time diagnosed anorexia nervosa and bulimia nervosa across 16 years in a danish nationwide psychiatric registry study. International Journal of Eating Disorders. 2015;48(7):845–50.

2. Smink FR, van Hoeken D, Hoek HW. Epidemiology of eating disorders: incidence, prevalence and mortality rates. Curr Psychiatry Rep. 2012;14(4):406–14.

3. 3. Washington APP. Diagnostic and statistical manual of mental disorders. DSM- 5, Fifth edition Edition.2013

4. Moskowitz L, Weiselberg E. Anorexia Nervosa/Atypical Anorexia Nervosa. Curr Probl Pediatr Adolesc Health Care. 2017;47(4):70–84.

5. Morita C, Tsuji H, Hata T, Gondo M, Takakura S, Kawai K, et al. Gut Dysbiosis in Patients with Anorexia Nervosa. PLoS One. 2015;10(12):e0145274.

6. Hanachi M, Manichanh C, Schoenenberger A, Pascal V, Levenez F, Cournède N, et al. Altered host-gut microbes symbiosis in severely malnourished anorexia nervosa (AN) patients undergoing enteral nutrition: An explicative factor of functional intestinal disorders? Clin Nutr. 2019;38(5):2304–10.

7. Di Lodovico L, Mondot S, Doré J, Mack I, Hanachi M, Gorwood P. Anorexia nervosa and gut microbiota: A systematic review and quantitative synthesis of pooled microbiological data. Prog Neuropsychopharmacol Biol Psychiatry. 2021;106:110114.

8. Mörkl S, Lackner S, Müller W, Gorkiewicz G, Kashofer K, Oberascher A, et al. Gut microbiota and body composition in anorexia nervosa inpatients in comparison to athletes, overweight, obese, and normal weight controls. Int J Eat Disord. 2017;50(12):1421–31.

9. Kleiman SC, Glenny EM, Bulik-Sullivan EC, Huh EY, Tsilimigras MCB, Fodor AA, et al. Daily Changes in Composition and Diversity of the Intestinal Microbiota in Patients with Anorexia Nervosa: A Series of Three Cases. European eating disorders review : the journal of the Eating Disorders Association. 2017;25(5):423–7.

10. Liang D, Leung RK, Guan W, Au WW. Involvement of gut microbiome in human health and disease: brief overview, knowledge gaps and research opportunities. Gut Pathog. 2018;10:3.

11. Fan Y, Støving RK, Berreira Ibraim S, Hyötyläinen T, Thirion F, Arora T, et al. The gut microbiota contributes to the pathogenesis of anorexia nervosa in humans and mice. Nature Microbiology. 2023;8(5):787–802.

12. Levy M, Kolodziejczyk AA, Thaiss CA, Elinav E. Dysbiosis and the immune system. Nature Reviews Immunology. 2017;17(4):219–32.

13. Mack I, Cuntz U, Grämer C, Niedermaier S, Pohl C, Schwiertz A, et al. Weight gain in anorexia nervosa does not ameliorate the faecal microbiota, branched chain fatty acid profiles, and gastrointestinal complaints. Sci Rep. 2016;6:26752.

14. Fouladi F, Bulik-Sullivan EC, Glenny EM, Thornton LM, Reed KK, Thomas S, et al. Reproducible changes in the anorexia nervosa gut microbiota following inpatient therapy remain distinct from non-eating disorder controls. Gut Microbes. 2022;14(1):2143217.

15. Schulz N, Belheouane M, Dahmen B, Ruan VA, Specht HE, Dempfle A, et al. Gut microbiota alteration in adolescent anorexia nervosa does not normalize with short-term weight restoration. Int J Eat Disord. 2021;54(6):969–80.

16. Prochazkova P, Roubalova R, Dvorak J, Tlaskalova-Hogenova H, Cermakova M, Tomasova P, et al. Microbiota, Microbial Metabolites, and Barrier Function in A Patient with Anorexia Nervosa after Fecal Microbiota Transplantation. Microorganisms. 2019;7(9).

17. de Clercq NC, Frissen MN, Davids M, Groen AK, Nieuwdorp M. Weight Gain after Fecal Microbiota Transplantation in a Patient with Recurrent Underweight following Clinical Recovery from Anorexia Nervosa. Psychother Psychosom. 2019:1–3.

18. Wilson BC, Derraik JGB, Albert BB, Leong KSW, Tweedie-Cullen RY, Creagh C, et al. An open-label pilot trial of faecal microbiome transfer to restore the gut microbiome in anorexia nervosa: protocol. BMJ Open. 2023;13(7):e070616.

19. Hata T, Miyata N, Takakura S, Yoshihara K, Asano Y, Kimura-Todani T, et al. The Gut Microbiome Derived From Anorexia Nervosa Patients Impairs Weight Gain and Behavioral Performance in Female Mice. Endocrinology. 2019;160(10):2441–52.

20. Glenny EM, Fouladi F, Thomas SA, Bulik-Sullivan EC, Tang Q, Djukic Z, et al. Gut microbial communities from patients with anorexia nervosa do not influence body weight in recipient germ-free mice. Gut Microbes. 2021;13(1):1–15.

21. Lundberg R. Humanizing the gut microbiota of mice: Opportunities and challenges. Lab Anim. 2019;53(3):244–51.

22. Lundberg R, Toft MF, August B, Hansen AK, Hansen CH. Antibiotic-treated versus germ-free rodents for microbiota transplantation studies. Gut Microbes. 2016;7(1):68–74.

23. Hansen AK, Krych Ł, Nielsen DS, Hansen CH. A Review of Applied Aspects of Dealing with Gut Microbiota Impact on Rodent Models. Ilar j. 2015;56(2):250–64.

24. Li X-m, Shi X, Yao Y, Shen Y-c, Wu X-l, Cai T, et al. Effects of Stool Sample Preservation Methods on Gut Microbiota Biodiversity: New Original Data and Systematic Review with Meta-Analysis. Microbiology Spectrum. 2023;11(3):e04297–22.

25. Gensollen T, Iyer SS, Kasper DL, Blumberg RS. How colonization by microbiota in early life shapes the immune system. Science. 2016;352(6285):539-44.

26. Méquinion M, Caron E, Zgheib S, Stievenard A, Zizzari P, Tolle V, et al. Physical activity: benefit or weakness in metabolic adaptations in a mouse model of chronic food restriction? American Journal of Physiology-Endocrinology and Metabolism. 2015;308(3):E241–E55.

27. Boekhorst J, Venlet N, Procházková N, Hansen ML, Lieberoth CB, Bahl MI, et al. Stool energy density is positively correlated to intestinal transit time and related to microbial enterotypes. Microbiome. 2022;10(1):223.

28. Rosenbaum M, Knight R, Leibel RL. The gut microbiota in human energy homeostasis and obesity. Trends Endocrinol Metab. 2015;26(9):493–501.

29. Fetissov SO. Role of the gut microbiota in host appetite control: bacterial growth to animal feeding behaviour. Nat Rev Endocrinol. 2017;13(1):11–25.

30. Beglinger C, Degen L. Gastrointestinal satiety signals in humans--physiologic roles for GLP-1 and PYY? Physiol Behav. 2006;89(4):460–4.

31. Houseknecht KL, Baile CA, Matteri RL, Spurlock ME. The biology of leptin: a review. J Anim Sci. 1998;76(5):1405–20.

32. Woods SC, Lutz TA, Geary N, Langhans W. Pancreatic signals controlling food intake; insulin, glucagon and amylin. Philosophical Transactions of the Royal Society B: Biological Sciences. 2006;361(1471):1219-35.

33. Misra M, Miller KK, Tsai P, Gallagher K, Lin A, Lee N, et al. Elevated peptide YY levels in adolescent girls with anorexia nervosa. J Clin Endocrinol Metab. 2006;91(3):1027–33.

34. Batterham RL, Cowley MA, Small CJ, Herzog H, Cohen MA, Dakin CL, et al. Gut hormone PYY(3-36) physiologically inhibits food intake. Nature. 2002;418(6898):650-4.

35. Stanley S, Wynne K, McGowan B, Bloom S. Hormonal Regulation of Food Intake. Physiological Reviews. 2005;85(4):1131–58.

36. Eckert ED, Pomeroy C, Raymond N, Kohler PF, Thuras P, Bowers CY. Leptin in Anorexia Nervosa1. The Journal of Clinical Endocrinology & Metabolism. 1998;83(3):791–5.

37. Grinspoon S, Gulick T, Askari H, Landt M, Lee K, Anderson E, et al. Serum leptin levels in women with anorexia nervosa. The Journal of Clinical Endocrinology & Metabolism. 1996;81(11):3861–3.

38. Hebebrand J, Muller TD, Holtkamp K, Herpertz-Dahlmann B. The role of leptin in anorexia nervosa: clinical implications. Molecular Psychiatry. 2007;12(1):23–35.

39. Van Harmelen V, Reynisdottir S, Eriksson P, Thörne A, Hoffstedt J, Lönnqvist F, et al. Leptin secretion from subcutaneous and visceral adipose tissue in women. Diabetes. 1998;47(6):913–7.

40. El Ghoch M, Calugi S, Lamburghini S, Dalle Grave R. Anorexia Nervosa and Body Fat Distribution: A Systematic Review. Nutrients. 2014;6(9):3895–912.

41. Wilcox G. Insulin and insulin resistance. Clin Biochem Rev. 2005;26(2):19–39.

42. Neary NM, Small CJ, Druce MR, Park AJ, Ellis SM, Semjonous NM, et al. Peptide YY3–36 and Glucagon-Like Peptide-17–36 Inhibit Food Intake Additively. Endocrinology. 2005;146(12):5120–7.

43. De Silva A, Salem V, Long CJ, Makwana A, Newbould RD, Rabiner EA, et al. The gut hormones PYY 3-36 and GLP-1 7-36 amide reduce food intake and modulate brain activity in appetite centers in humans. Cell Metab. 2011;14(5):700–6.

44. Schmidt JB, Gregersen NT, Pedersen SD, Arentoft JL, Ritz C, Schwartz TW, et al. Effects of PYY3–36 and GLP-1 on energy intake, energy expenditure, and appetite in overweight men. American Journal of Physiology-Endocrinology and Metabolism. 2014;306(11):E1248–E56.

45. Seitz J, Dahmen B, Keller L, Herpertz-Dahlmann B. Gut Feelings: How Microbiota Might Impact the Development and Course of Anorexia Nervosa. Nutrients. 2020;12(11).

46. Kameyama K, Itoh K. Intestinal Colonization by a Lachnospiraceae Bacterium Contributes to the Development of Diabetes in Obese Mice. Microbes and Environments. 2014;29(4):427–30.

47. Liu D, Wen B, Zhu K, Luo Y, Li J, Li Y, et al. Antibiotics-induced perturbations in gut microbial diversity influence metabolic phenotypes in a murine model of high-fat diet-induced obesity. Applied Microbiology and Biotechnology. 2019;103(13):5269–83.

48. Roager HM, Licht TR. Microbial tryptophan catabolites in health and disease. Nat Commun. 2018;9(1):3294.

49. Sanmiguel CP, Jacobs J, Gupta A, Ju T, Stains J, Coveleskie K, et al. Surgically Induced Changes in Gut Microbiome and Hedonic Eating as Related to Weight Loss: Preliminary Findings in Obese Women Undergoing Bariatric Surgery. Psychosom Med. 2017;79(8):880–7.

50. Borgo F, Riva A, Benetti A, Casiraghi MC, Bertelli S, Garbossa S, et al. Microbiota in anorexia nervosa: The triangle between bacterial species, metabolites and psychological tests. PLoS One. 2017;12(6):e0179739.

51. Rizk M, Mattar L, Kern L, Berthoz S, Duclos J, Viltart O, et al. Physical Activity in Eating Disorders: A Systematic Review. Nutrients. 2020;12(1).

52. Miletta MC, Iyilikci O, Shanabrough M, Šestan-Peša M, Cammisa A, Zeiss CJ, et al. AgRP neurons control compulsive exercise and survival in an activity-based anorexia model. Nature Metabolism. 2020;2(11):1204–11.

53. Poffé C, Dalle S, Kainz H, Berardi E, Hespel P. A noninterfering system to measure in-cage spontaneous physical activity in mice. Journal of Applied Physiology. 2018;125(2):263–70.

54. Mähler Convenor M, Berard M, Feinstein R, Gallagher A, Illgen-Wilcke B, Pritchett-Corning K, et al. FELASA recommendations for the health monitoring of mouse, rat, hamster, guinea pig and rabbit colonies in breeding and experimental units. Lab Anim. 2014;48(3):178–92.

55. Smith AJ, Clutton RE, Lilley E, Hansen KEA, Brattelid T. PREPARE: guidelines for planning animal research and testing. Lab Anim. 2018;52(2):135–41.

56. Percie du Sert N, Hurst V, Ahluwalia A, Alam S, Avey MT, Baker M, et al. The ARRIVE guidelines 2.0: Updated guidelines for reporting animal research. PLoS Biol. 2020;18(7):e3000410.

57. Percie du Sert N, Ahluwalia A, Alam S, Avey MT, Baker M, Browne WJ, et al. Reporting animal research: Explanation and elaboration for the ARRIVE guidelines 2.0. PLoS Biol. 2020;18(7):e3000411.

58. Ellekilde M, Selfjord E, Larsen CS, Jakesevic M, Rune I, Tranberg B, et al. Transfer of gut microbiota from lean and obese mice to antibiotic-treated mice. Scientific Reports. 2014;4(1):5922.

59. Ubeda C, Taur Y, Jenq RR, Equinda MJ, Son T, Samstein M, et al. Vancomycin-resistant Enterococcus domination of intestinal microbiota is enabled by antibiotic treatment in mice and precedes bloodstream invasion in humans. The Journal of Clinical Investigation. 2010;120(12):4332–41.

60. Zachariassen LF, Sørensen DB, Krych L, Hansen AK, Hansen CHF. Effects of delivery mode on behavior in mouse offspring. Physiol Behav. 2021;230:113285.

61. Andersen-Civil AIS, Myhill LJ, Büdeyri Gökgöz N, Engström MT, Mejer H, Zhu L, et al. Dietary proanthocyanidins promote localized antioxidant responses in porcine pulmonary and gastrointestinal tissues during Ascaris suum-induced type 2 inflammation. The FASEB Journal. 2022;36(4):e22256.

62. Arildsen AW, Zachariassen LF, Krych L, Hansen AK, Hansen CHF. Delayed Gut Colonization Shapes Future Allergic Responses in a Murine Model of Atopic Dermatitis. Frontiers in Immunology. 2021;12.

63. Quast C, Pruesse E, Yilmaz P, Gerken J, Schweer T, Yarza P, et al. The SILVA ribosomal RNA gene database project: improved data processing and web-based tools. Nucleic Acids Research. 2012;41(D1):D590–D6.

