## Supplementary Table 1,2, Supplementary Figures 1 - 5 for "Investigating the Rescue Potential of Fecal Microbiota Transplants in Anorexia Nervosa using Antibiotic-treated Mice in a Split Group Cross-Over Study"

**Supplementary Material**

Supplementary Table 1. Comparison of accumulated food intake between donor groups after FMT1 and FMT2. Overall, the AN cages after FMT1 and ANAN cages after FMT2 had lower food intake between 1.9% and 6.9% accumulated in any comparison with other donor combinations.

| Treatment | % Less Food | Exp | FMT | Duration | Age of mice | Cages | Number of mice |
|---|---|---|---|---|---|---|---|
| AN vs CO | 6.89% | A | FMT1 | 5 | 10-11 weeks | 6 | 24 |
| AN vs CO | 1.85% | B | FMT1 | 5 | 10-11 weeks | 6 | 24 |
| AN/AN vs CO/CO | 6.25% | A | FMT2 | 4 | 16-17 weeks | 6 | 12 |
| AN/AN vs CO/CO | 6.53% | B | FMT2 | 4 | 16-17 weeks | 6 | 12 |
| AN/AN vs AN/CO | 2.19% | A | FMT2 | 4 | 16-17 weeks | 6 | 12 |
| AN/AN vs CO/AN | 5.43% | A | FMT2 | 4 | 16-17 weeks | 6 | 12 |
| AN/AN vs AN/CO | 4.12% | B | FMT2 | 4 | 16-17 weeks | 6 | 12 |
| AN/AN vs CO/AN | 2.97% | B | FMT2 | 4 | 16-17 weeks | 6 | 12 |

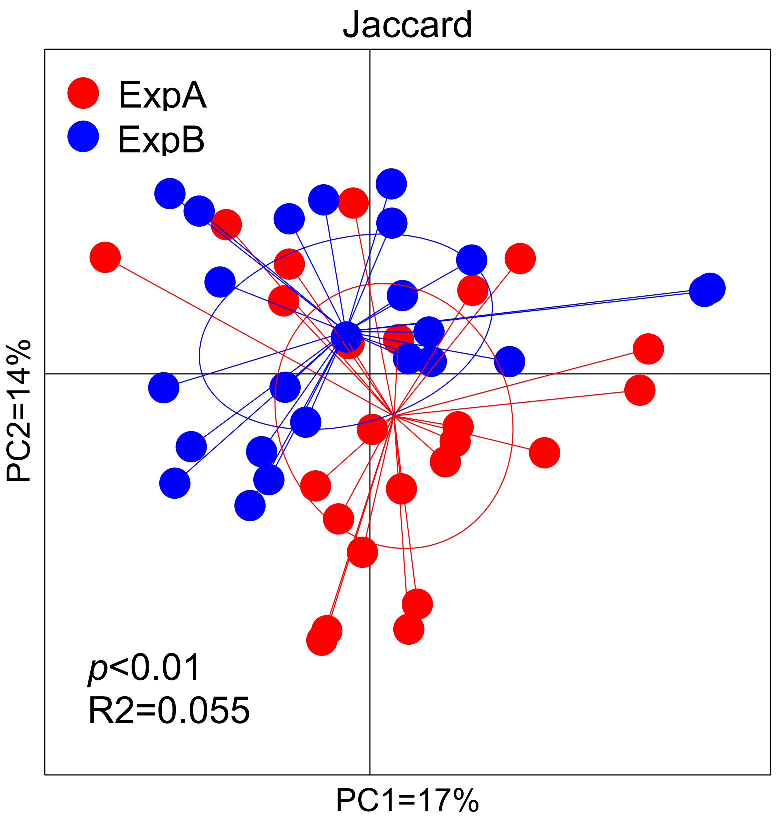


Supplementary Figure 1. Differences of gut microbiota after FMT1 between the experimental runs ExpA and ExpB shown by the Jaccard index. After FMT1 the two runs were statistically significant different when looking at the presence and absence of ASVs (**$p < 0.01$).

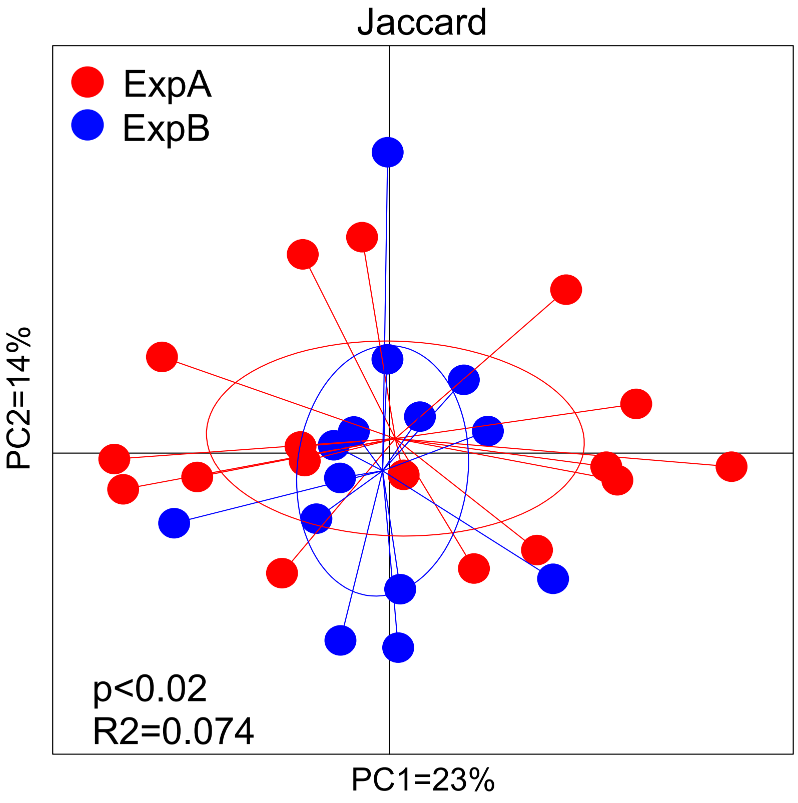


Supplementary Figure 2. Differences of gut microbiota after FMT2 between the experimental runs ExpA and ExpB shown by the Jaccard index. After FMT2 the two runs were statistically significant different when looking at the presence and absence of ASVs (*$p < 0.02$).

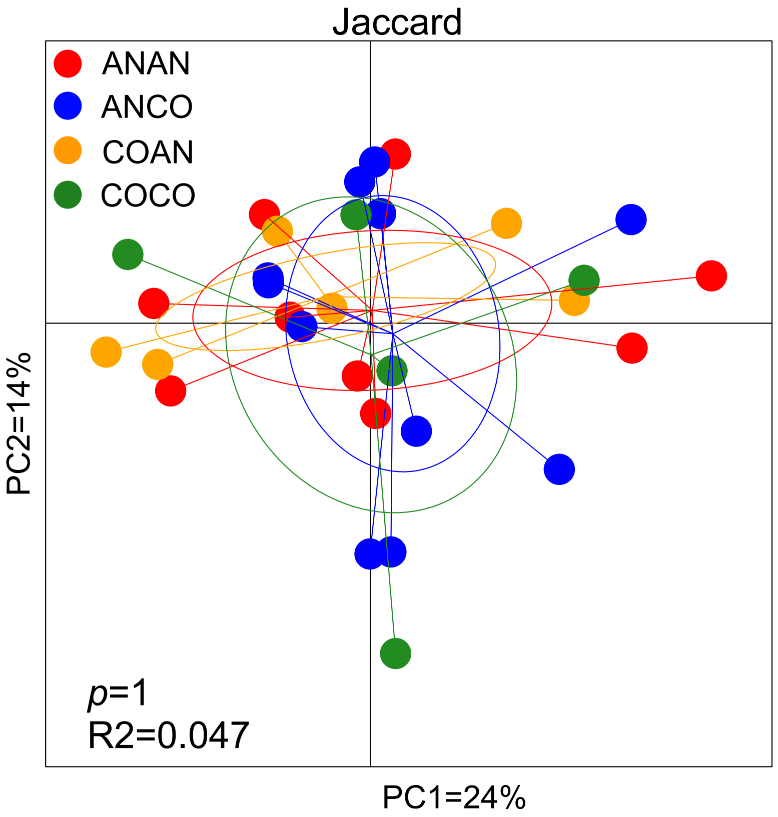


Supplementary Figure 3. After two weeks of AB treatment and four weeks of FMT2, the combined donor types did not differ significantly from each other (ANAN = red, ANCO = blue, COAN = yellow, COCO = green).

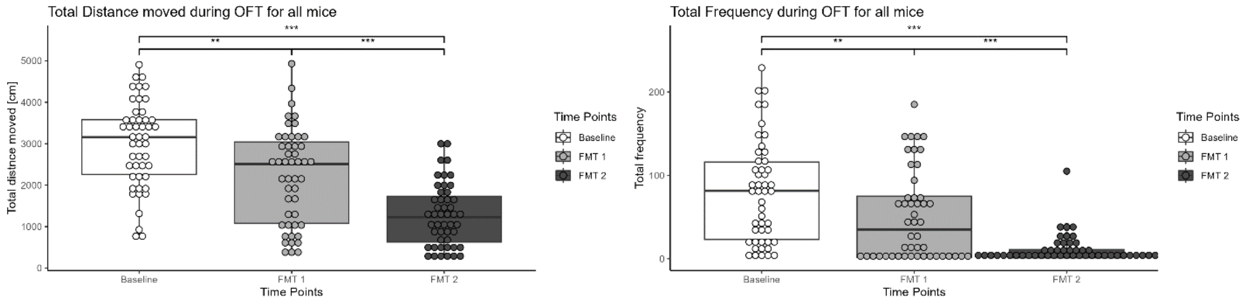


Supplementary figure 4. Effects of time, treatments and repeated measurements on general locomotor activity and total frequency. All mice (n = 48) were subjected to open field test before FMT1 (**Baseline**), after FMT1 (**FMT1**), and after FMT2 (**FMT2**). General locomotor activity and total frequency was evaluated by paired t-test (2-tailed). **A)** There was significant decrease in general locomotor activity after FMT1 (****p* < 0.001) and after FMT2 (****p* < 0.0001). **B)** There was significant difference in total frequency after FMT1 (***p* < 0.01) and after FMT2 (****p* < 0.0001).

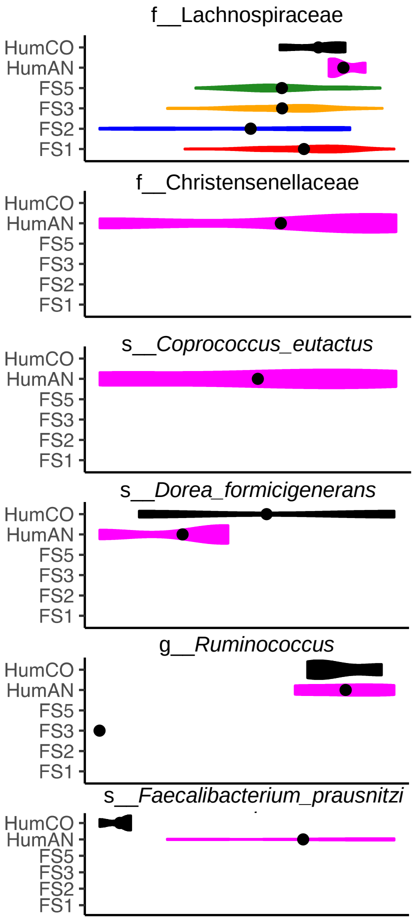


Supplementary Figure 5. Significant difference in bacterial compositions between the mice fecal samples obtained at various time points and the human donors. *Lachnospiraceae* fluctuates from different time points and is more abundant in the human AN samples compared to the CO samples (top graph). There is a difference in abundance of *Dorea formicigenerans* between human AN and CO samples. The abundance of *Faecalibacterium prausnitzi* is higher in human AN samples compared to CO samples.

Supplementary table 2. Overview of primers used to target the hypervariable region (V1-V8) when sequencing near full-length 16S rRNA gene amplicons using the oxford nanopore technology (ONT).

| Name | Sequence |
|---|---|
| UMI_338Fa | GTCTCGTGGG CTCGGNNNNN NNNNNNNNNN ACWCCTACGG GWGGCAGCAG |
| UMI_338Fb | GTCTCGTGGG CTCGGNNNNN NNNNNNNNNN GACTCCTACG GGAGGCWGCA G |

| | |
|---|---|
| UMI_1391R | GTCTCGTGGG CTCGGNNNNN NNNNNNNNNN GACGGGCGGT GTGTRCA |
| UMI_27Fa | GTCTCGTGGG CTCGGNNNNN NNNNNNNNNN AGAGTTTGAT YMTGGCTYAG |
| UMI_27Fb | GTCTCGTGGG CTCGGNNNNN NNNNNNNNNN AGGGTTCGAT TCTGGCTCAG |
| UMI_1540_R | GTCTCGTGGG CTCGGNNNNN NNNNNNNNNN TACGGYTACC TTGTTACGAC T |
